# LHX2 regulates dendritic morphogenesis in layer II/III of the neocortex via distinct pathways in progenitors and postmitotic neurons

**DOI:** 10.1101/2024.01.30.577728

**Authors:** Mahima Bose, Sreenath Ravindran, Sanjna Kumari, Achintya Srivastava, Archana Iyer, Binita Vedak, Ishita Talwar, Rishikesh Narayanan, Shubha Tole

**Author notes:** equal contributors (Mahima Bose, Sreenath Ravindran).

## Abstract

In the mammalian neocortex, excitatory neurons that send projections via the corpus callosum are critical to integrating information across the two brain hemispheres. The molecular mechanisms governing the development of the dendritic arbours and spines of these callosal neurons are poorly understood, yet these features are critical to their physiological properties. LIM Homeodomain 2 (*Lhx2*), a regulator of fundamental processes in cortical development, is expressed in postmitotic callosal neurons occupying layer II/III of the neocortex and also in their progenitors residing in the embryonic day (E) 15.5 ventricular zone of the mouse neocortex. We tested whether this factor is essential for dendritic arbour configuration and spine morphogenesis of layer II/III neurons. Here, we report loss of *Lhx2,* either in postmitotic layer II/III neurons or their progenitors, resulted in shrunken dendritic arbours and perturbed spine morphology. The defects were more pronounced upon *Lhx2* disruption in progenitors, and were recapitulated when this was driven exclusively in basal progenitors. In postmitotic neurons, LHX2 regulates dendritic and spine morphogenesis via the canonical Wnt /β Catenin signalling pathway. Constitutive activation of this pathway in postmitotic neurons mimics the *Lhx2* loss-of-function phenotype. In E15.5 progenitors, LHX2 acts in part via bHLH transcription factor NEUROG2 to regulate dendritic morphogenesis. We demonstrate that loss of *Lhx2* causes a massive increase in *Neurog2* expression, and that *Neurog2* knockdown partially rescues the loss of *Lhx2* phenotype. Our study uncovers novel LHX2 functions consistent with its temporally dynamic and diverse roles in development.

**Teaser:** The mature architecture of a neuron is shaped by distinct genetic mechanisms that act in its mother cell and after it is born.

## Introduction

The functional characteristics of a neuron are critically dependent on its dendritic arborisation and complexity (*1*). Disruptions of dendritic morphogenesis underlie neurodevelopmental disorders, including Autism Spectrum Disorder and schizophrenia (*2*, *3*). In the mammalian cerebral cortex, superficial pyramidal layer neurons, which occupy layers II/III, broadly extend their apical dendrites into layer I and basal dendrites within their layer. They also establish connections with the contralateral hemisphere via the corpus callosum. These neurons are born late in cortical neurogenesis, with their production peaking at embryonic day (E) 15.5 in the mouse (*4*).

Dendritic morphogenesis is regulated by multiple factors, including transcriptional regulators, guidance cues, cell adhesion molecules, cytoskeletal elements, and neuronal activity (*5*, *6*). Among these, transcription factors are known to be the drivers of downstream signalling pathways and gene expression responsible for dendritic development (*7*). In the developing mammalian cerebral cortex, transcription factors like *Cux1*, *Cux2*, *Zfp312*, and *Foxg1* are known to promote dendritic arborisation in specific neuronal subtypes, and the loss of any of these genes leads to reduced complexity (*8–12*). On the other hand, factors like *Sox11, Tbr1* and *Neurog2* have been shown to suppress excessive branching, such that the loss of these factors leads to increased complexity (*13–15*).

Despite the established knowledge regarding the involvement of various transcription factors in regulating dendritogenesis, the precise stage of differentiation and maturation at which their role is relevant remains to be determined. Some factors, *e.g. Zfp312, Sox11*, *Tbr1,* are known to regulate morphometry exclusively in specific subtypes of postmitotic neurons (*11*, *14*, *15*), while others, such as *Foxg1,* display a broader expression pattern encompassing different progenitors and neuronal classes (*12*, *16*). This complexity prompts an additional question: At which stage of neuronal maturation is dendritic morphometry determined? Is the modulation of dendritic architecture solely a postmitotic event, or does a mechanism for regulating the dendritic structure of a neuron exist in its progenitor? The temporal dynamics of transcription factor engagement in regulating dendritogenesis during the different stages of development, *i.e.*, progenitor and postmitotic, remain poorly explored.

LHX2 is a pleiotropic transcription factor implicated in multiple facets of cortical development, including cortical patterning (*17–21*), progenitor proliferation (*22–24*), cell fate decisions (*25*) and neuron subtype specification (*26*, *27*). Its expression in the cortical primordium is seen as early as E8.5 and is an established regulator of distinct stages of cortical development (*28*). LHX2 function in early cortical progenitors is critical for properly developing the thalamocortical tract and barrel formation (*24*, *27*) and producing the correct numbers of layer VI versus layer V neurons (*26*, *29*). In postmitotic layer IV cortical neurons, *Lhx2* is essential for the activity-dependent orientation of dendrites towards the barrel fields (*30*). A broad cortex-specific loss of *Lhx2* results in a lack of the corpus callosum and hippocampal commissure (*31*). These distinct functions in cortical progenitors and postmitotic neurons motivated the question of whether this factor may regulate a particular process, that of dendritic arborisation and spine maturation, at multiple stages of neuronal differentiation and whether it utilises independent pathways to do so at each stage.

Here, we show that loss of *Lhx2,* either in cortical progenitors or specifically in postmitotic neurons, causes profound alterations of the dendritic and spine morphology of layer II/III neurons while sparing corpus callosum crossing. We demonstrate that the progenitor-specific role is recapitulated by loss of *Lhx2* restricted to basal progenitors. We identify downstream target NEUROG2 as an effector of LHX2 function by utilising shRNA approaches to show that simultaneous knockdown of *Neurog2* can partially rescue the dendritic arborisation phenotype resulting from loss of *Lhx2.* In postmitotic neurons, we report that LHX2 suppresses the Wnt/β-catenin signalling cascade. We demonstrate that independent stabilisation of β-catenin in postmitotic neurons recapitulates the effect of loss of *Lhx2* on dendritic arborisation and spine morphology.

In summary, our findings reveal novel insights into the distinct cell-stage-specific roles of transcription factor LHX2 in orchestrating diverse gene networks that govern the dendritic and spine morphogenesis process.

## Results

Layer II/III cortical neuronal birth peaks at embryonic day (E) 15.5 in mice (*4*). LHX2 protein is detected in the cortical progenitors at this stage and persists in postmitotic layer II/III neurons (Figure 1A, B). This observation motivated us to compare the effects of loss of *Lhx2,* either in E15.5 progenitors or in newborn postmitotic neurons, on the dendritic morphogenesis of mature layer II/III neurons. All dendritic arbour and spine morphology measurements were performed on postnatal day (P) 30 in the somatosensory cortex layer II/III. To label the dendritic arbours, we used *in utero* electroporation (IUE) to introduce a plasmid expressing membrane-bound EGFP (CAAX-EGFP) in E15.5 progenitors so that the neurons arising from these cells were labelled in perpetuity (Figure 1C, C’). We used an *Lhx2 ^lox/lox^* mouse line to effect conditional loss of *Lhx2*. Postmitotic neuron-specific disruption of *Lhx2* was effected by crossing this line to a line expressing *Neurod6-Cre (NexCre;* (*32*). IUE of CAAX-EGFP at E15.5 produced EGFP-labelled layer II/III neurons, which then experienced *Lhx2* gene disruption upon becoming postmitotic due to the action of *NexCre*.

**Figure 1:**
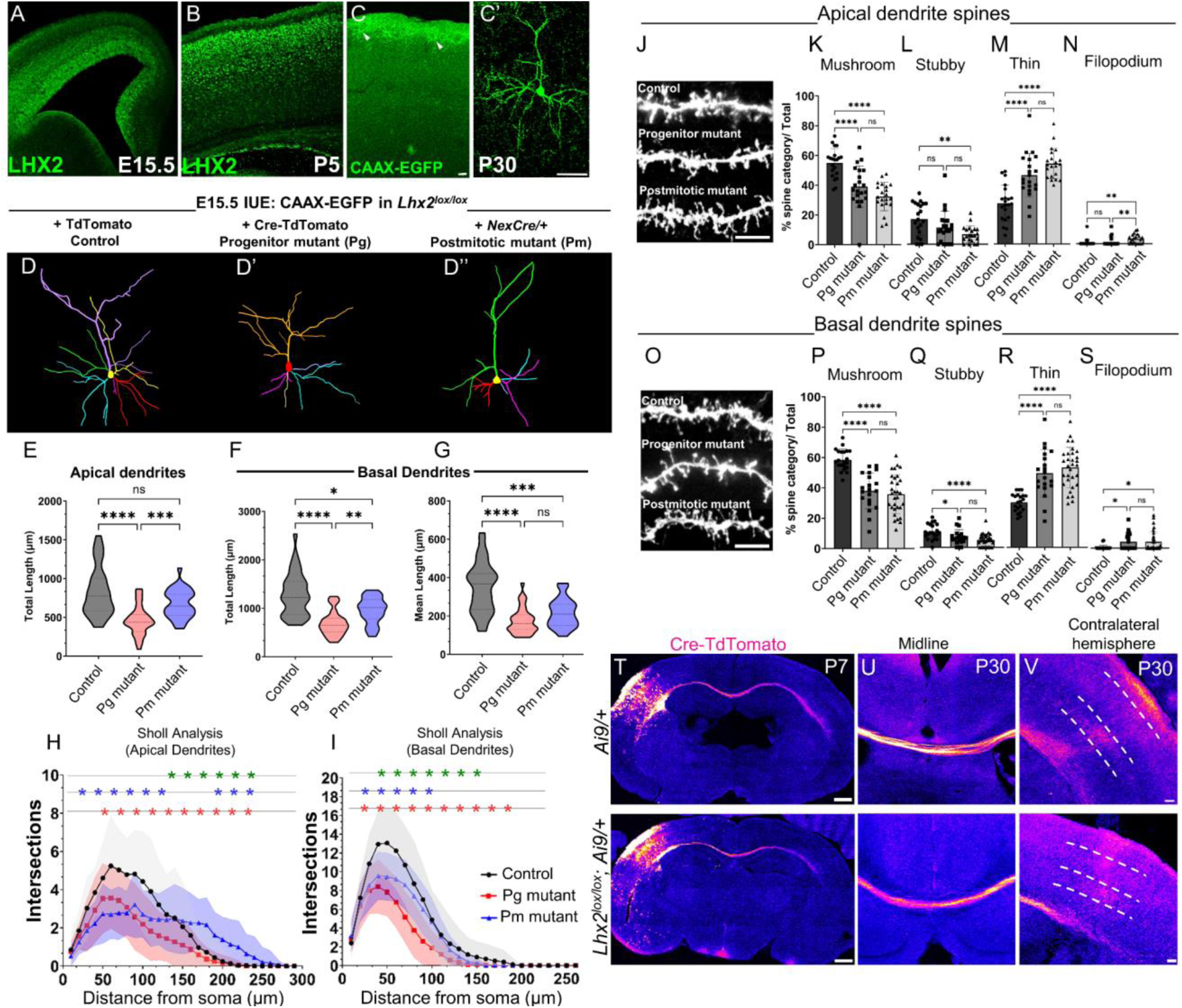
Loss of *Lhx2* either in progenitors or newborn postmitotic neurons induces dendritic morphology and dendritic spine defects in layer II/III neocortical neurons. (A, B) LHX2 in progenitors at E15.5 (A) and layer II/III cortical neurons at P5 (B). (C, C’) A P30 layer II/III neuron labelled by CAAX-EGFP IUE in E15.5 progenitors (D–D’’) Neurolucida tracings of Control, Progenitor (Pg), and Postmitotic (Pm) *Lhx2* mutant neurons at P30. (E–G) The total length of apical dendrites (E) is significantly reduced in Pg mutant neurons (Control: 831.3 ± 316.4μm, Pg mutant: 460.2 ± 185μm; Pm mutant: 677 ± 181.9μm). The mean length (F) and total length (G) of basal dendrites (F) are significantly reduced for Pg and Pm mutant neurons (F: Control: 348.6 ± 131.2μm; Pg mutant: 174.0 ± 66.7μm; Pm mutant: 215.2 ± 74.4μm); (G: Control: 1285 ± 438.7μm; Pg mutant: 687 ± 249.7μm; Pm mutant: 956 ± 265.4μm). (H, I) Sholl analysis of apical dendrites (H) and basal dendrites (I) shows a significant decrease in complexity in Pg and Pm mutants. (35 to 40 neurons were scored for each condition obtained from 4 biologically independent replicates; *Statistical test: Two-way ANOVA-multiple comparisons with Tukey’s correction for Sholl analysis and Kruskal-Wallis Test for total/mean length analysis; asterisks indicate pairwise comparisons between Control vs Pg mutant (red), Control vs Pm mutant (blue) and Pg mutant vs Pm mutant (green). detailed statistical significance values per intersection row provided in Supplementary Figure S5 and Data S1*). (J–S) Dendritic spine morphology is affected in Pg and Pm mutant neurons. In apical and basal dendrites, the percentage of spines with mushroom morphology was significantly reduced (K, P) with a concomitant increase in thin morphology (M, R). The percentage of stubby spines was also reduced in apical dendrites of Pm mutants (L) and basal dendrites of Pg and Pm mutants (Q). The percentage of filopodium spines increased in Pm apical dendrites (N) and Pg basal dendrites (S). (20 dendrites with 25 spines each were analysed from each category. *Statistical Test: Kruskal-Wallis Test*). (T–V) Axons of Control and *Lhx2* Pg mutant neurons are seen in the corpus callosum at P7 (T) and arborise in the contralateral hemisphere by P30 (U, V). *\*(p<0.05), **(p<0.01), ***(p<0.001), ****(p<0.0001). Scale bars in (C, C’, T, V) are 50 μm and in (J, O) are 5 μm*.

To disrupt *Lhx2* in progenitors, we co-electroporated a plasmid encoding Cre-TdTomato with CAAX-EGFP at E15.5 (Supplementary Figure S1G–I). As controls, TdTomato (lacking Cre) was used.

### Disrupting *Lhx2* in E15.5 progenitors and newly post-mitotic neurons causes dendritic and spine morphology defects in P30 layer II/III neurons

To analyse the fully developed dendritic arborisation of layer II/III neurons, we examined CAAX-EGFP-labelled cells at P30 when dendritic morphogenesis is complete. Control neurons displayed a typical layer II/III pyramidal morphology (Figure 1C’, D).

In contrast, both progenitor-specific (Pg; Fig. 1D’) and postmitotic (Pm; Fig. 1D’’) displayed shrunken dendritic arbours upon loss of *Lhx2*. The total length of apical and basal dendrites was significantly reduced in Pg and Pm mutants, with a more pronounced effect seen in Pg mutants than in Pm mutants. The mean length of the basal dendrites exhibited a severe reduction in both Pg and Pm perturbations (Figure 1E–G). Sholl analysis of apical and basal dendrites reflected decreased branching complexity in both Pg and Pm mutant neurons.This effect was stronger in Pg mutants than in Pm mutants in both apical and basal dendrites (Figure 1H, I).

We tested whether loss of *Lhx2* in progenitors affects the subtype identity of the mutant layer II/III neurons since we had previously reported a switch in subtype identity from layer VI to layer V when *Lhx2* was disrupted in deep layer progenitors (*26*). However, preliminary analysis showed that Pg mutant neurons expressed the upper layer marker POU3F2 similar to controls (Supplementary Figure S1A–F) and displayed a characteristic pyramidal morphology (Supplementary Figure S1G–I), suggesting that neuronal subtype identity is broadly preserved.

These results collectively suggest that *Lhx2* plays a crucial role in regulating the dendritic morphogenesis of both apical and basal dendrites in layer II/III neurons, operating in progenitors and postmitotic neurons.

### Disrupting *Lhx2* in E15.5 progenitors and postmitotic neurons causes spine development defects in layer II/III neurons

Dendritic spines serve as the functional units of dendrites and are the sites of synaptic transduction (*33*, *34*). Understandably, the molecular factors influencing dendritic development overlap with those involved in spine development (*34–36*). We investigated whether the loss of *Lhx2* in either progenitor or postmitotic neurons affects the formation of spines in P30 layer II/III neuronal dendrites. Control, Pg, and Pm mutant neurons were analysed for spine morphology in apical and basal dendrites. Spines were categorised into filopodial, thin, stubby, and mushroom types, which correlates with the overall maturation stage of the spines (*34*). We observed a significant increase in the percentage of thin spines and a corresponding decrease in mushroom spines in the apical and basal dendrites of both Pg and Pm mutants. Stubby spines were significantly fewer in Pm but not Pg mutant apical dendrites (Figure 1L). These findings indicate a role for LHX2 in the morphogenesis of spines during neurodevelopment to achieve the mushroom shape typical of mature spines.

### Corpus Callosum crossing is unaffected upon loss of *Lhx2*

Axons originating from neurons in layer II/III of the neocortex play a crucial role in corpus callosum formation (*37*, *38*). Previous studies reported that loss of *Lhx2* driven by *Emx1-*Cre or *Nestin-*Cre, which act in cortical progenitors from E11.5, results in corpus callosum agenesis, although inactivation using *Nex-Cre* did not (*31*). We sought to investigate whether the impact of *Lhx2* on corpus callosum development is cell-autonomous by electroporating Cre-TdTomato in the progenitors of *Lhx2^lox/lox^; Ai9/+* (mutant) and *Ai9/+* (Control) mice at E15.5. The corpus callosum was examined at P7 and P30.

In contrast to the striking effect on Layer II/III dendrites, the axons of these neurons did not appear to be significantly affected in the mutants. The corpus callosum traversed the midline (Figure 1T–V), reached the contralateral hemisphere (Figure 1T), and developed arbours at layer V and layer II/III by P30 (Figure 1V). This finding conclusively shows that the corpus callosum agenesis reported using *Emx1*-Cre or *Nestin*-Cre (*31*) is not due to a cell-autonomous role in the neurons of layer II/III and motivates exploration of additional roles for *Lhx2* in midline guidance mechanisms as suggested in (*31*).

In summary, although LHX2 is a pleiotropic factor, its function in the maturation of layer II/III neurons is confined to their dendritic processes and spines, not their axons. Furthermore, the loss of *Lhx2* at the Pg stage appeared to have a stronger effect on dendritic morphology than at the Pm stage. In the following sections, we examined the molecular mechanisms by which LHX2 regulates these processes in postmitotic layer II/III neurons and their progenitors.

### Loss of *Lhx2* increases the firing rate and excitability of layer II/III neurons

We sought to examine the consequences of the morphometric deficits resulting from the loss of *Lhx2* on the electrophysiological properties of neocortical layer II/III pyramidal neurons. We measured several sub-and supra-threshold intrinsic physiological properties from brain slices of Pg and Pm *Lhx2* mutant neurons using patch-clamp electrophysiology and compared them with their respective controls (Figure 2, Supplementary Figures S3–S7). Dendritic arborisation is a well-established factor in regulating neuronal excitability (*39*, *40*), therefore we first measured input resistance, a subthreshold measure of neuronal gain. Consistent with the more pronounced reduction in dendritic arbour in Pg *Lhx2* mutant neurons (Figure 1), we found a significant increase in input resistance values in this condition compared to their control values (Figure 2A, B). However, control and Pm *Lhx2* mutant neurons manifested comparable input resistance values (Figure 2C, D). Consistently, impedance amplitude, another measure of sub-threshold excitability, was significantly higher than controls in Pg mutant neurons but were comparable to their controls in Pm mutant neurons (Supplementary Figures S3–S4). Other physiological measurements of sub-threshold excitability were comparable in Pg and Pm mutant neurons compared to their respective controls (Supplementary Figures S3–S4).

**Figure 2:**
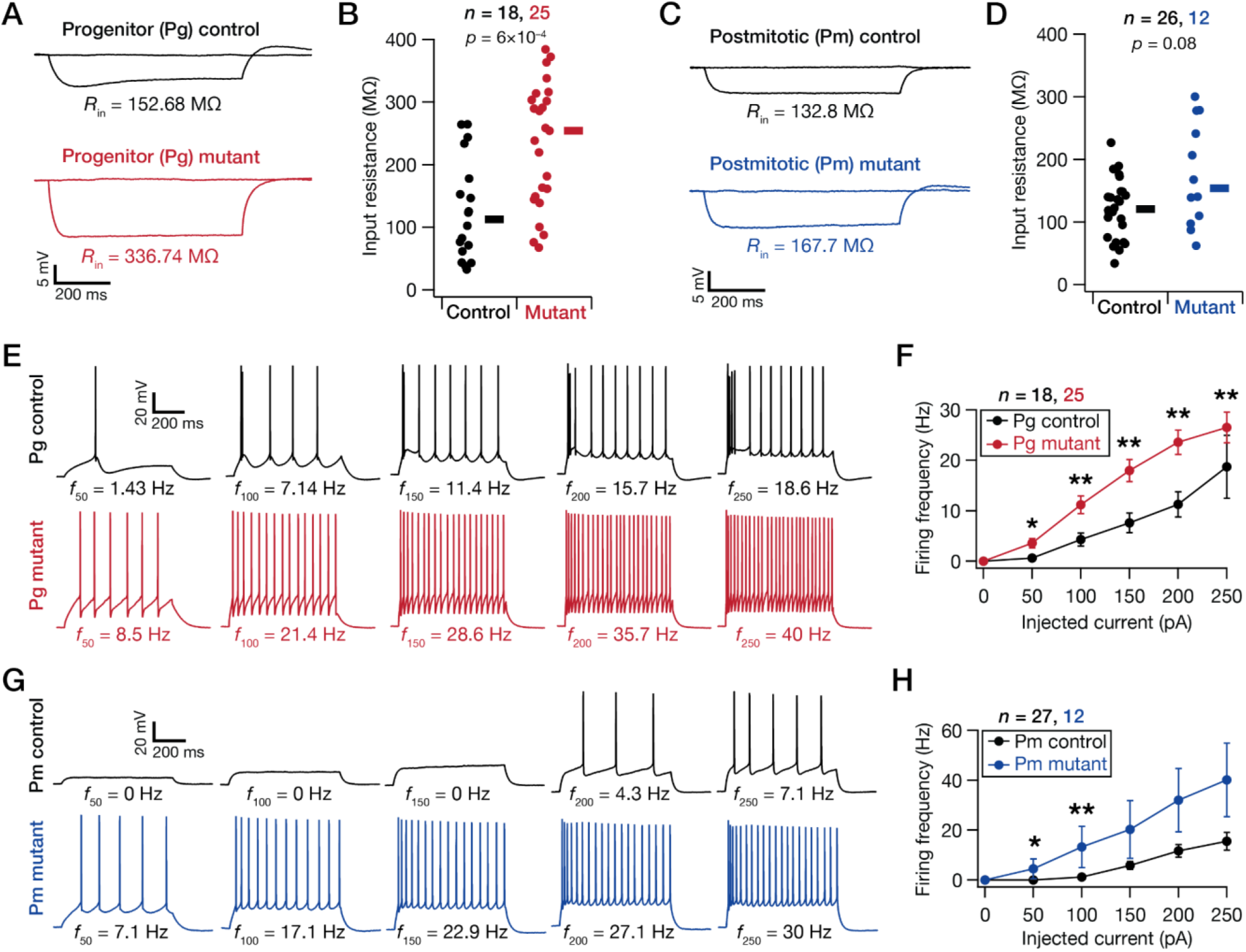
Loss of *Lhx2* either in progenitors (Pg) or newborn postmitotic (Pm) neurons enhanced intrinsic excitability of neocortical layer II/III pyramidal neurons. (A) Example voltage responses of neurons to pulse current injections of –50 pA and 0 pA, used for computing input resistance (*R*_*in*_). (B) Beeswarm plots of input resistance (*R*_*in*_) obtained from control and Pg mutant neurons. The thick line on the right of each beeswarm plot depicts respective median values. (C, D) Same as panels *A–B*, for Pm mutant neurons and associated controls. (E) Voltage traces recorded from example pyramidal neurons in response to 50–250 pA pulse current injections from control (top panels) and Pg mutant (bottom panels) mice. The firing rate associated with each trace is provided at the bottom of the trace. (F) Mean-SEM plot showing firing rates of neurons from control and Pg mutant mice, plotted against current injection amplitudes. (G, H) Same as panels E–F, for Pm mutant neurons and associated controls. *Statistical test: Wilcoxon rank sum test: *(p<0.05), **(p<0.01). n represents the number of total neurons recorded in each group.* Additional analyses of sub-and supra-threshold measurements from all four groups are provided in Supplementary Figs. S3–S7.

In striking contrast, we found both Pg (Figure 2E, F) and Pm mutant neurons (Figure 2G, H) to manifest significantly higher supra-threshold excitability than in their respective controls. Specifically, we found significantly higher firing frequencies (Figure 2E–H, Supplementary Figure S5) and a switch to depolarisation-induced block with low current injections (Supplementary Figure S5C, D). In addition, Pm mutant neurons displayed a significant reduction in both the latency to the first spike (Supplementary Figure S6), the first inter-spike interval (Supplementary Figure S6) and a significant increase in the peak *dV*/*dt* associated with the action potentials (Supplementary Figure S6). These measurements were not significantly different between Pg mutant neurons and their controls (Supplementary Figure S7).

In summary, loss of *Lhx2* at the Pg or Pm stage resulted in distinct effects on specific electrophysiological measurements, but in both conditions there was significant enhancement in the action potential firing rate in layer II/III neocortical pyramidal neurons (Figure 2, Supplementary Figures S3–S7).

### LHX2 regulates dendritic development in postmitotic neurons by suppressing WNT/ β Catenin signalling cascade genes

Since the postmitotic loss of *Lhx2* was driven by *NexCre,* which acts in the entire postmitotic cortical plate, we tested whether the role of LHX2 is cell autonomous using prepared primary dissociated cultures. Embryos from control and *NexCre/+; Lhx2^lox/lox^* embryos were first electroporated *ex vivo* with CAAX-EGFP at E15.5 for labelling purposes and then dissociated and maintained *in vitro* for 15 days. We observed a significant decrease in the total length of both apical and basal dendrites, similar to the phenotype *in vivo* (Fig. 3A–D), suggesting that the function of LHX2 in regulating dendritic development is cell-autonomous.

**Figure 3:**
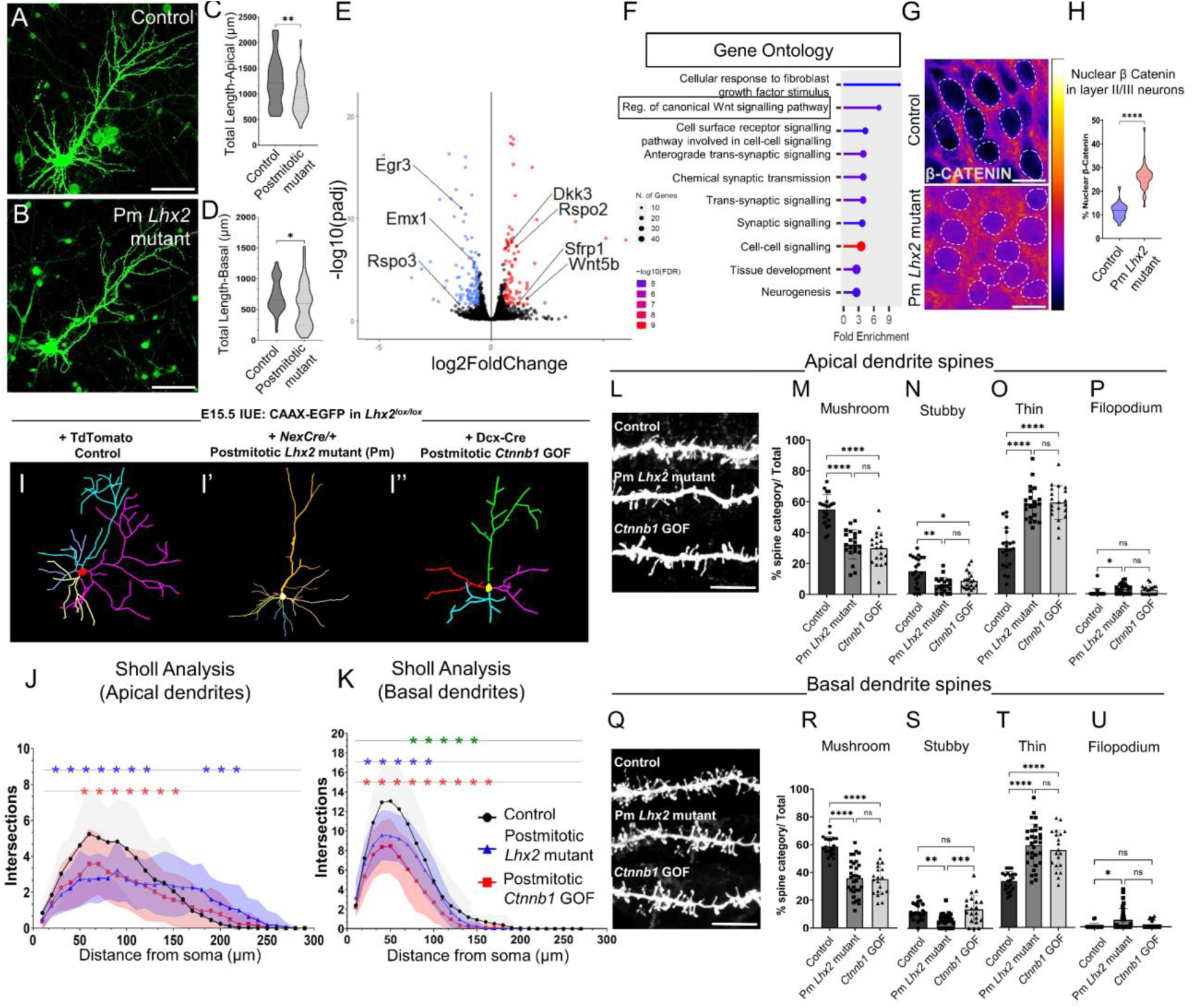
*Lhx2* regulates canonical Wnt signalling in postmitotic neurons. (A–D) *In vitro* neuronal cultures (A, B) display a reduction in the total length of apical (C) and basal (D) dendrites in Pm *Lhx2* mutant neurons. (E, F) RNA Sequencing of control and postmitotic mutant neurons at P5 (E) reveals dysregulated Wnt signalling cascade genes, as confirmed by Gene Ontology (GO) analysis (F). (G, H) Layer II/III Pm *Lhx2* mutant neurons display increased nuclear localisation of the Wnt signalling downstream effector, β CATENIN. (I, I’’) Neurolucida tracings of control, Pm *Lhx2* mutant, and Doublecortin-Cre-driven *Ctnnb1* gain-of-function (GOF) neurons at P30. (J, K) Sholl analysis of apical and basal dendrites shows similar dendritic defects in Pm *Lhx2* mutant and *Ctnnb1* GOF neurons (30–40 neurons in each category were analysed from 4 biologically independent experiments. *Statistical test: Two-way ANOVA-multiple comparisons with Tukey’s correction; asterisks indicate pairwise comparisons between Control vs Ctnnb1* GOF *(red), Control vs Pm mutant (blue) and Pm mutant vs Ctnnb1* GOF *(green) detailed statistical significance values per intersection row provided in Supplementary Figure S8 and Data S1)*. (L–U) Dendritic spine morphology defects are comparable between Pm *Lhx2* mutant and *Ctnnb1* GOF neurons. Both apical and basal dendrites exhibit a significant reduction in mushroom and stubby morphology spines (M, R) and a concomitant increase in thin and filopodium morphology spines (O, T; .20 dendrites with 25 spines each were analysed for each category. *Statistical test: Kruskal-Wallis Test for spine morphology analysis*)*. *(p<0.05), **(p<0.01), ***(p<0.001), ****(p<0.0001). Scale bars are 50 μm in (A, B), 10μm in (G) and 5 μm in (L, Q)*.

To gain insights into the mechanistic role of LHX2 in postmitotic neurons, we isolated control and Pm mutant neurons at P5, when the dendritic arbours are maturing, and performed RNA Sequencing. As before, the neurons were labelled by *in utero* electroporation of CAAX-EGFP in *NexCre/+, Lhx2^loxlox^* brains and littermate controls at E15.5. RNA Seq analysis revealed that several differentially expressed genes were associated with the Wnt signalling pathway, such as *Dkk3, Rspo2, Sfrp1*, and *Wnt5b* (Figure 3E). This was further confirmed in the Gene Ontology analysis of differentially expressed genes (Figure 3F). A typical readout of active canonical Wnt signalling is the stabilisation of cytoplasmic β-Catenin and its translocation to the nucleus (*41*). We validated this by immunostaining for β-CATENIN and discovered a significant increase in its nuclear localisation in layer II/III Pm mutant neurons (Figure 3G, H). These findings suggested that postmitotic loss of *Lhx2* enhances canonical Wnt signalling in layer II/III neurons.

To ascertain whether upregulation of canonical Wnt signalling can regulate dendritic morphogenesis, we examined whether directly stabilising β-catenin can recapitulate the dendritic dendritic defects observed upon loss of *Lhx2*.

We used a *Ctnnb1* (β-CATENIN) conditional gain-of-function (GOF) mouse line in which exon 3 is floxed. Loss of this exon prevents β-CATENIN from being phosphorylated and tagged for degradation, thus resulting in a constitutively stabilised β-CATENIN protein (*42*). To effect loss of *Ctnnb1* specifically in postmitotic neurons, we electroporated a plasmid encoding Doublecortin (*Dcx*)-driven Cre recombinase together with CAAX-EGFP for labelling purposes. We analysed the outcomes at P30, focusing on EGFP-labelled neurons born at E15.5 from *Dcx*-Cre+CAAX-EGFP electroporated progenitors, which would be expected to display β-catenin GOF upon becoming postmitotic.

Sholl analysis of the apical dendrites of the *Ctnnb1* GOF neurons showed a striking recapitulation of the apical and basal dendritic defects seen in postmitotic *Lhx2* mutant condition (Fig. 3I–I’’, J, K). *Ctnnb1* GOF neurons also recapitulated the spine defects observed in postmitotic *Lhx2* mutant neurons, *i.e.* an increase in the proportion of thin morphology spines and a concomitant decrease in mushroom morphology spines in both apical and basal dendrites (Fig. 3L–U). In summary, these findings indicate that LHX2 suppresses Wnt/ β Catenin signalling in postmitotic layer II/III neurons and that this may be the critical pathway via which LHX2 mediates the regulation of dendritic and spine morphogenesis in these cells.

Next, we tested whether LHX2 recruits act via similar or distinct mechanisms in progenitors to regulate these processes.

### Loss of *Lhx2* in progenitors results in an upregulation in *Neurog2* at postnatal stages

Although progenitor-specific loss of *Lhx2* using IUE already tests cell-autonomous functions due to the sparse nature of the electroporation we employed, we sought to ascertain whether *Lhx2* mutant progenitors can recapitulate the *in vivo* dendritic arborisation phenotype (Figure 1) in neurons that are born *in vitro* (Fig. 4A, B). We prepared dissociated cultures from *Lhx2^lox/lox^* cortices electroporated *ex vivo* with either TdTomato+CAAX-EGFP (control) or Cre-TdTomato+CAAX-EGFP (Progenitor mutant) at E15.5 and cultured the cells *in vitro* for 15 days. Remarkably, the progenitor mutant neurons exhibited similar phenotypes *in vitro*, significantly reducing the total lengths of apical and basal dendrites (Fig. 4C, D).

**Figure 4:**
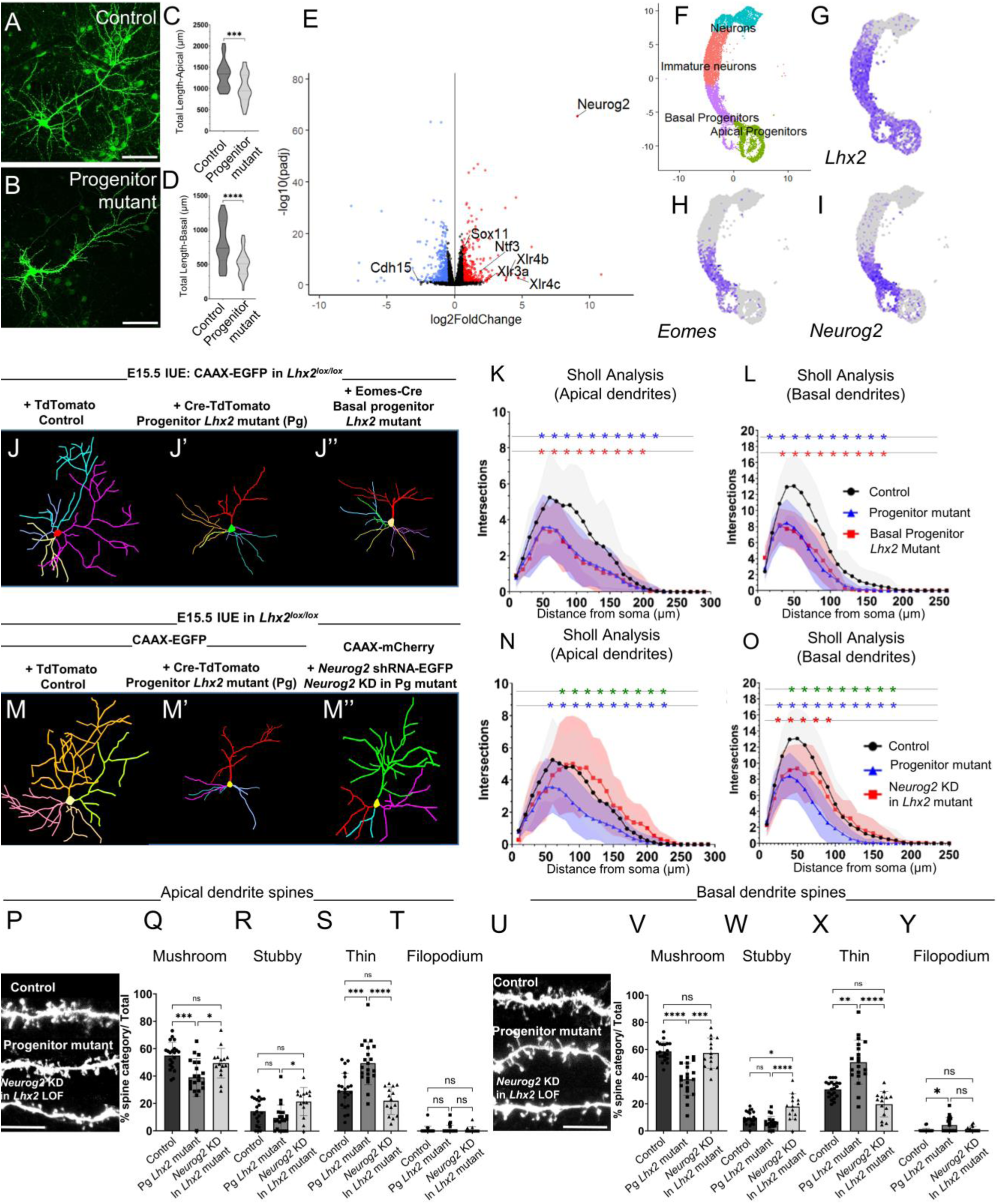
LHX2 modulates dendritic morphogenesis of neurons by suppression of *Neurog2* in progenitors. (A–D) *In vitro* neuronal cultures display a reduction in the total length of apical (C) and basal (D) dendrites in Pg *Lhx2* mutant neurons. (E) RNA Sequencing of control and Pg *Lhx2* mutant neurons at P5 shows significant upregulation of *Neurog2*. (F–I) Analysis of E15.5 single-cell RNA seq data shows *Lhx2* expression in apical, basal progenitors and immature neurons (G), while *Neurog2* expression (I) specifically in *Eomes*-positive (H) basal progenitors. (J–L) Dendritic morphology analysis of Control, Pg *Lhx2* mutant neurons and *pEomes-*Cre driven basal progenitor *Lhx2* mutant layer II/III neurons. (J–J’’) Neurolucida tracings and Sholl analysis of apical (K) and basal (L) dendrites reveal that loss of *Lhx2* in basal progenitors recapitulates the pan-progenitor Pg *Lhx2* mutant phenotype. (M–O) Dendritic morphology analysis of Control, Pg *Lhx2* mutant neurons, and Pg *Lhx2* mutant neurons with *Neurog2* shRNA (M–M’’) Neurolucida tracings and Sholl analysis reveals that *Neurog2* knockdown in Pg *Lhx2* mutant neurons leads to a significant and complete rescue in apical dendrites (N) and a significant partial rescue in basal dendrites (O). (30–35 neurons were analysed from 3 biologically independent experiments. *Statistical test: Two-way ANOVA-multiple comparisons with Tukey correction; asterisks in (K, L) indicate pairwise comparisons between Control vs Basal progenitor mutant (red), Control vs Pg mutant (blue) and Pg mutant vs basal progenitor mutant (green); asterisks in (N,O) indicate individual comparisons between Control vs Neurog2 KD in Lhx2 mutant (red), control vs Pg mutant (blue) and Pg mutant vs Neurog2 KD in Lhx2 mutant (green); detailed statistical significance values per intersection row provided in Supplementary Figure S8 and Data S1)*. (P–Y) Spine morphology analysis of apical and basal dendrites of Control, Pg and *Neurog2* KD in *Lhx2* Pg mutant neurons reveals a significant amelioration of spine morphology defects. *\*(p<0.05), **(p<0.01), ***(p<0.001), ****(p<0.0001). Scale bars are 50 μm in (A, B) and 5 μm in (P, U)*.

**Figure 5:**
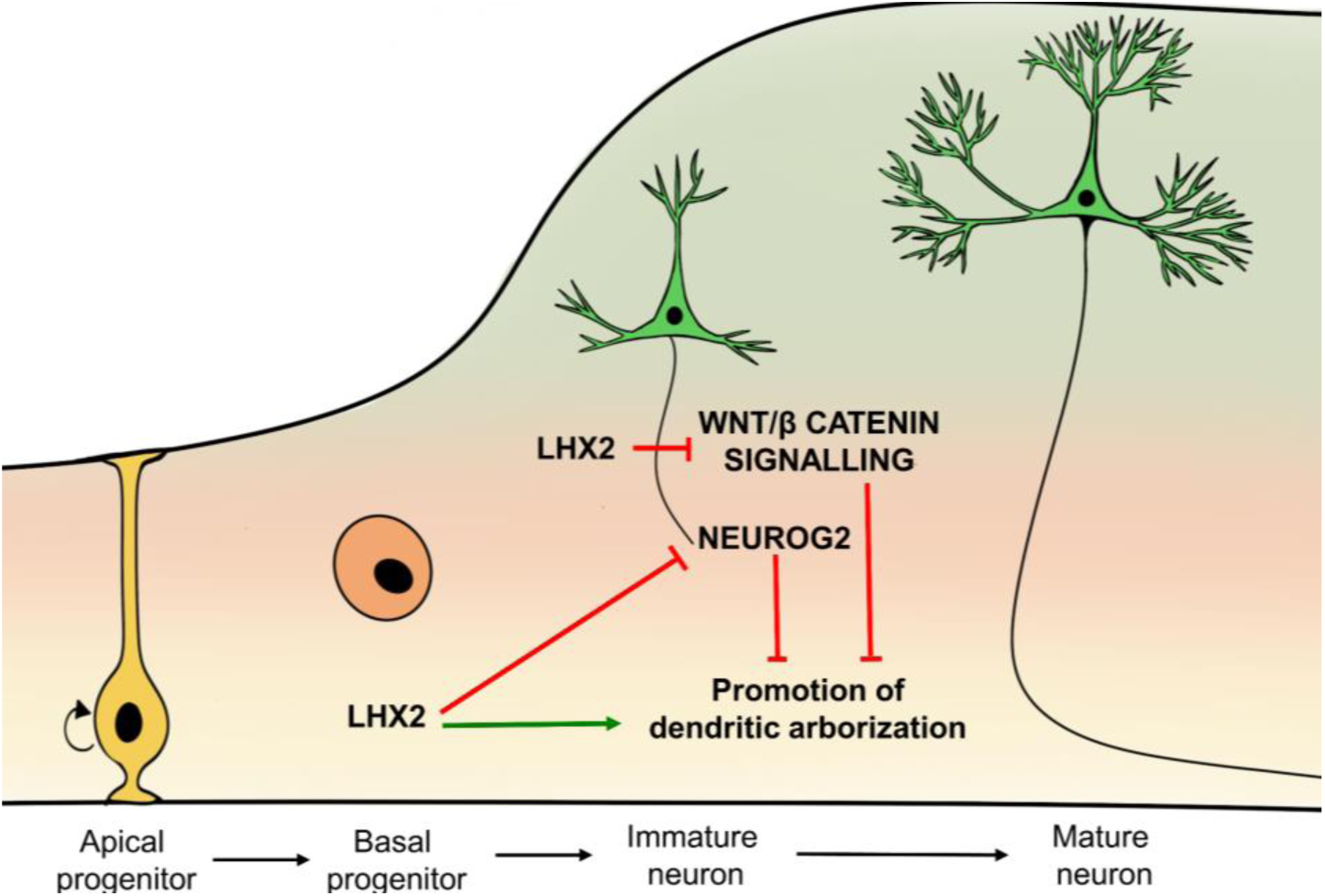
Schematic depicting the mechanism of action of LHX2 during the different stages of neuron development. LHX2 plays distinct roles in regulating gene networks during dendritic morphogenesis in progenitors and postmitotic neurons. LHX2 function in progenitors controls *Neurog2* expression in postmitotic neurons, facilitating its downregulation as cells differentiate from basal progenitors. Moreover, in postmitotic neurons, LHX2 regulates the Wnt/β-catenin signalling pathway by repressing active Wnt/β Catenin signalling cascade genes, thereby promoting proper dendritic arborisation of layer II/III cortical neurons.

To explore the molecular basis of LHX2 function, we isolated E15.5-electroporated control and progenitor mutant neurons at P5 and performed RNA Seq (Figure 4E). Loss of *Lhx2* resulted in dysregulation of numerous genes governing dendrite and spine development, including *Ntf3, Xlr4b, Xlr3b, Xlr4c, Sox11*, and *Cdh15* (*8*, *9*, *14*, *43*). The top dysregulated gene was transcription factor *Neurog2,* which was 535-fold upregulated in the *Lhx2* mutant neurons (Figure 4E). *Neurog2* expression was unchanged in Pm *Lhx2* mutant neurons (Figure 3E), indicating that this is a Pg-specific requirement of *Lhx2* that manifests at later stages.

### The progenitor mutant defect is recapitulated upon loss of *Lhx2* exclusively in basal progenitors

NEUROG2 is a transcriptional regulator previously implicated in neuronal differentiation, migration, and morphogenesis (*13*, *44*). Analysis of the single-cell RNA Seq dataset of E15.5 neocortical cells obtained from (*45*) revealed that while *Lhx2* is enriched in both apical and basal progenitors during neurogenesis, *Neurog2* is specifically enriched in the *Eomes*-positive basal progenitors but is downregulated in postmitotic neurons (Figure 4F–I; (*46*)). This led us to investigate whether the function of *Lhx2* in regulating dendritic morphogenesis is specific to basal, and not apical progenitors. To test this, we utilised an *Eomes*-driven Cre recombinase (*47*) to effect disruption of *Lhx2* specifically in basal progenitors (Fig. 4J–J’’).

Sholl analysis of apical and basal dendrites of *Eomes*-Cre driven *Lhx2* mutant neurons demonstrated a near-perfect overlap of the two progenitor mutant graphs (Cre-and *Eomes*-Cre), indicating that the role of LHX2 in regulating dendrite morphometry is relevant specifically in basal progenitors that go on to produce layer II/III neurons (Figure 4K–L).

### Knockdown of *Neurog2* partially rescues the *Lhx2* progenitor mutant dendritic defect

Having identified *Neurog2* to be significantly upregulated upon progenitor-specific loss of *Lhx2*, we investigated whether downregulation of *Neurog2* in the *Lhx2* mutant progenitors could rescue the dendritic and spine morphology defects. To test this, we employed a short-hairpin RNA construct to knock down *Neurog2 (*(*48*); Supplementary Figure S11) in the progenitors together with Cre and CAAX-mCherry in the *Lhx2^lox/lox^* background, creating a simultaneous knockout of the *Lhx2* gene and a knockdown of *Neurog2* expression. Sholl analysis of these neurons at P30 demonstrated a significant and complete rescue of the dendritic defect phenotype in the apical dendrites and a significant partial rescue in the basal dendrites (Figure 4M–O). Spine morphology analysis in the apical and basal dendrites of these neurons also revealed substantial amelioration of the spine defects observed in Pg *Lhx2* mutants (Figure 4P–Y). In contrast, knockdown of *Neurog2* alone resulted in increased complexity of apical dendrites and no change in basal dendrites (Supplementary Figure S12).

In summary, these results indicate that LHX2 plays a crucial role in E15.5 basal progenitors which results in suppression of *Neurog2* expression by the postnatal stage. This is required for the normal development of dendritic arbours and spine morphogenesis in layer II/III cortical neurons.

## Discussion

Our results reveal distinct roles of LHX2 in cortical progenitors and postmitotic neurons that govern unique gene regulatory networks critical for dendritic and spine morphometry and the electrophysiological properties of neurons.

We identify an LHX2-NEUROG2 interaction that modulates dendritic morphometry. NEUROG2 is a well-established factor essential in the differentiation, migration, and maturation of neurons (*13*, *44*, *49*, *50*). Our findings extend these functions to the modulation of dendritic morphometry and spine maturation. The Pg mutant paradigm we use (electroporation of Cre or basal progenitor-specific (*Eomes*-Cre) induces a long term loss of *Lhx2* that also affects postmitotic neurons arising from the mutant progenitors and also results in a massive upregulation of *Neurog2* in these cells. This paradigm does not allow an assessment of the stage at which *Neurog2* upregulation is relevant in terms of functional consequences. However, the fact that it is not normally expressed in postmitotic neurons, but is present only at earlier stages (*46*), is suggestive that relevant stage of *Neurog2* dysregulation may the postmitotic neuronal stage. The upregulation of *Neurog2* in postmitotic neurons is not seen if *Lhx2* is disrupted exclusively at postmitotic stages. Therefore, we have uncovered a unique progenitor-specific function of LHX2 that results in *Neurog2* upregulation subsequently.

Loss of *Lhx2* and downregulation of *Neurog2* individually have opposite effects on apical dendritic complexity, which is decreased in the former and increased in the latter condition.

When performed together, there is a complete normalization of apical dendritic complexity. These results indicate that in addition to *Neurog2* being a target of LHX2, each factor may have opposite effects on common or distinct downstream targets that regulate apical dendritic complexity. In the case of basal dendrites, downregulation of *Neurog2* alone has no effect, and loss of *Lhx2* alone reduces complexity. When performed together, there is a partial rescue of basal dendritic complexity. These results indicate that in addition to acting via *Neurog2,* LHX2 may act via distinct targets the dysregulation of which is not normalized by the concurrent knockdown of *Neurog2* and loss of *Lhx2.* Since NEUROG2 is a pleiotropic transcriptional regulator (*51*) implicated in pioneering chromatin interactions essential for differentiation (*49*), and LHX2 regulates its expression during neurogenesis, understanding how these factors function in tandem to regulate specific developmental programs will be a rich line of future investigation.

These findings also offer novel insights into how LHX2 adapts its roles at different stages of neurogenesis. Previous studies identified an early cortical progenitor-specific role of LHX2 that regulates the electrophysiological properties of the firstborn cells of the cortex, the subplate (*27*). These early (E11.5) progenitors give rise to deep-layer progenitors, in which LHX2 regulates subtype identity via control of layer V fate determinant *Fezf2* (*26*); the deep layer progenitors then give rise to superficial layer progenitors in which the current study reveals yet another role of LHX2, that of regulating dendritic morphogenesis of layer II/III neurons. Together, these studies demonstrate that the role of LHX2 in cortical neurogenesis evolves as progenitors progress through time, possibly due to temporally unique occupancy patterns of the factor on the genome, resulting in differences in gene regulation. This understanding offers potential for future studies, including a nuanced analysis of such temporally dynamic functions of a single transcription factor.

In Layer II/III postmitotic neurons, we identify that LHX2 regulates Wnt/β Catenin signalling. This pathway is well-established in regulating neuron polarity and dendritogenesis (*51*, *52*). However, our study adds a layer of complexity by showing that enhanced Wnt signalling in postmitotic layer II/III neurons of the lateral somatosensory cortex reduces dendritic complexity, contrary to its function in the medial cortex (*43*, *53*). LHX2, in turn, regulates active Wnt signalling in these postmitotic neurons, contributing to proper dendritogenesis and circuit maturation. This provides insight into region-specific differences in the functionality and responsiveness of signalling cascades and their regulation.

In the developing telencephalon, the cortical hem is a localised medial source of Wnt ligands, and medial cortical structures such as the hippocampus and the retrosplenial cortex display corresponding responsiveness in upregulating target genes (*42*, *43*, *54*). Consequently, the role of Wnts in dendritogenesis in these structures has been well studied and shown to have a positive growth response to enhanced Wnt signalling (*43*, *53*, *55*). However, other members of the Wnt ligand family, such as *Wnt7b,* are expressed widely in the dorsal telencephalon (*56*). This raises the question of whether all cortical regions in the telencephalon interpret Wnt signals similarly, resulting in the same developmental outcomes. Our findings suggest a complex scenario in which enhanced Wnt signalling in postmitotic layer II/III neurons of the lateral somatosensory cortex reduces dendritic complexity, whereas, an opposite role has been reported in the medial cortex (*43*, *53*, *55*). The unique molecular context of each neuronal subtype likely influences the overall functional outcomes downstream of the same pathway. LHX2 suppresses Wnt signalling in lateral cortical postmitotic neurons, contributing to proper dendritogenesis and spine maturation, but its role in this process in the medial cortex has not yet been examined. Furthermore, since *Lhx2* is expressed in superficial layer neurons into adulthood, the possibility of a maintenance function for this factor, which may operate via the Wnt pathway in a regional-specific manner, needs to be examined. Together, these findings provide valuable insights into spatial differences in the functionality and responsiveness of signalling cascades and their transcriptional regulation.

We report that loss of *Lhx2* has profound consequences not only on the dendritic arbour of layer II/III neurons but also on their action potential firing properties, resulting in a pronounced increase in intrinsic excitability in mutant mice. These findings motivate new lines of inquiry into how LHX2 regulates specific biophysical properties of neurons.

In summary, our study offers novel insights into cell-type-specific and region-specific gene regulatory networks modulated by LHX2, contributing to neuronal maturation, function, and circuit development. LHX2 haploinsufficiency in humans leads to neurodevelopmental disorders, including autism spectrum disorder, intellectual disability and microcephaly (*57*). In this context, our findings provide a biological foundation for the molecular logic of how this pleiotropic transcription factor affects multiple processes in distinct stages and structures during development. This is an essential step in uncovering the basis of such neurodevelopmental disorders.

## Acknowledgements

We thank Dr Shital Suryavanshi and the animal house staff of the Tata Institute of Fundamental Research (TIFR) for their excellent support; Edwin Monuki for the *Lhx2* floxed mouse line; M. Mark Taketo (Kyoto University, Japan) for the *Ctnnb1* exon-3 floxed line; Dr Mahendra Sonawane (Tata Institute of Fundamental Research, India) for the CAAX-*Egfp* and CAAX-*mCherry* plasmids; Dr Tarik Haydar (University of Maryland, USA) for the p*Eomes*-Cre plasmid; and Dr François Guillemot (Francis Crick Institute, London, UK) for the pSilcaggs-*Neurog2*-shRNA-EGFP and pSilcaggs-scrambled-EGFP plasmids. We thank Medgenome Inc. for sequencing the RNA libraries. We thank Varun Suresh for his assistance with the initial processing of the RNA Sequencing data. This work was funded by the Department of Atomic Energy (DAE), Govt. of India (Project Identification no. RTI4003, DAE OM no. 1303/2/2019/R&D-II/DAE/2079).

## Methods

### Mice

All procedures followed the Tata Institute of Fundamental Research Animal Ethics Committee (TIFR-IAEC) guidelines. The *Lhx2^lox/lox^* mouse line was a gift from Edwin Monuki (University of California, Irvine, USA). The *Ai9* reporter mouse line (Strain #:007909) was obtained from Jackson Laboratory. The *NexCre/+* mouse line was received from Klaus Nave, Max Planck Institute for Experimental Medicine (*32*). The *Ctnnb1* conditional GOF mouse line was obtained from M. Mark Taketo (Kyoto University, Japan). All animals were kept at an ambient temperature and humidity, with a 12h light-dark cycle and food available ad libitum. Noon of the day of the vaginal plug was designated as embryonic day 0.5 (E0.5). Both male and female animals were used for all experiments.

All the electrophysiology recording experiments were reviewed and approved by the Institute Animal Ethics Committee of the Indian Institute of Science, Bangalore. Experimental procedures were similar to previously established protocols (*58–60*) and are detailed below. 7 (4 males and 3 females) mice from the Pm control group between 7–9 months old, 7 (3 males and 4 females) from Pm *Lhx2* mutant mice between 6-10 months old, 4 (2 males and 2 females) mice belonging to the Pg control group between 2–4 months old, and 5 (3 males and 2 females) Pg *Lhx2* mutant mice between 2-4 months old were used for *in vitro* patch-clamp electrophysiology experiments. Animals were provided *ad libitum* food and water and were housed with an automated 12-hour light-dark cycle, with the facility temperature maintained at 23° C.

Primers for genotyping were (expected band sizes):

*Lhx2* conditional forward: ACCGGTGGAGGAAGACTTTT

*Lhx2* conditional reverse: CAGCGGTTAAGTATTGGGACA

(WT: 144 bp, *Lhx2^lox^*: 188bp, Null: 231 bp)

*NexCre* Forward: GAGTCCTGGAATCAGTCTTTTTC

*NexCre* Reverse: AGAATGTGGAGTAGGGTGAC

*NexCre* Mutant Reverse: CCGCATAACCAGTGAAACAG

(WT: 770 bp, *NexCre*: 525 bp).

### *In utero* electroporation and DNA constructs

*In utero* electroporation was performed at E15.5 as previously described (*27*). Embryos were injected with plasmid DNA solution dissolved in nuclease-free water with 0.1% fast green with plasmid DNA into the lateral ventricle through the uterine wall using a fine glass microcapillary (Sutter capillaries #B100-75-10).

The pCAGGS-IRES-TdTomato construct was obtained from Addgene (#83029). pCAGGS-IRES-Cre-TdTomato was generated in the lab. Membrane-bound EGFP (CAAX-EGFP) and mCherry (CAAX-mCherry) were gifts from Dr Mahendra Sonawane (Tata Institute of Fundamental Research, India). p*Eomes*-Cre was obtained from Dr Tarik Haydar (University of Maryland, Baltimore, USA). p*Dcx*-Cre was previously described in (*61*). pSilcaggs-*Neurog2* shRNA-EGFP and pSilcaggs-scrambled-EGFP were gifts from François Guillemot (Francis Crick Institute, London, UK).

### Tissue Preparation

Electroporated mice were harvested on postnatal day 30. Mice were anaesthetised using Thiopentone and transcardially perfused with Saline, followed by 4% (wt/vol) paraformaldehyde in phosphate buffer. Brains were kept for overnight fixation and then cryoprotected by transferring to 30% sucrose-PBS until sectioning. The brains were sectioned at 40μm using a Vibratome (Leica VT2000S).

### Immunohistochemistry

Brains were sectioned at 40um and processed for free-floating immunohistochemistry. Sections were washed and permeabilised with phosphate buffer containing 0.3% (vol/vol) Triton X-100. Blocking was done with 5% (vol/vol) horse serum in phosphate buffer with 0.3% (vol/vol) Triton X-100 for one hour at room temperature. Primary antibody treatment was performed in phosphate buffer containing 0.3% (vol/vol) Triton X-100 with 2.5% (vol/vol) horse serum and incubated overnight at 4°C. The sections were washed in phosphate buffer, followed by the appropriate secondary antibody for two hours at room temperature and DAPI. Sections were mounted on Superfrost plus glass microscope slides (Cat #71869-10) and protected with Fluoroshield (Sigma cat no F6057 or F6182).

Antibodies used: Biotinylated GFP (Abcam; ab6658, 1:200), Rabbit RFP (Abcam; ab65856, 1:200), Neurog2 (ThermoFisher Scientific; PA5-78556, 1:1000), β Catenin (BD Biosciences; A610151, 1:200).

### Image acquisition and analysis

Images of neurons and dendritic spines were imaged using Olympus FluoView 1200 and 3000 confocal microscopes with FluoView software. A step size of 0.8um was kept to maximise the z-axis resolution of the dendritic branches. All the image analysis was done on Fiji-ImageJ. A nonlinear operation such as gamma correction was not performed in the images. Brightness and contrast adjustments were performed identically for control and mutant conditions.

### Dendritic morphology and spine analysis

3D morphology reconstruction of CAAX-EGFP/CAAX-mCherry labelled neurons was done using the Neurolucida 2017 software. The apical and basal dendrites were identified based on the directionality of the dendrites. The primary dendrite directed towards the pial surface of the cortex was considered apical, and the rest were labelled as basal. Total length, mean length and Sholl analysis were performed using the Neurolucida Explorer 2017 software. For dendritic spine analysis, high-resolution images of the distal branches of the apical and basal dendrites were acquired on the confocal system. The ImageJ plugin – Dendritic Spine Counter (https://imagej.net/plugins/dendritic-spine-counter) was used to classify spines into mushroom, stubby, thin or filopodium by calculating the neck length, neck width and head width of the spines. Approximately 15-20 dendrites with ∼25 spines each, obtained from 4 biological replicates, were analysed. Kruskal Wallis Test was performed to compare the three conditions. All statistical analysis was performed using GraphPad Prism 10.0.0 software.

### RNA Sequencing and analysis

For RNA Sequencing, the E15.5 electroporated layer II/III (Control: EGFP in *Lhx2^lox/lox^*, Progenitor mutant: Cre-EGFP in *Lhx2^lox/lox^*; Postmitotic mutant EGFP in *NexCre/+ Lhx2^lox/lox^*) was microdissected from the cortical tissue at P5 after sectioning on a tissue chopper. RNA extraction and sequencing were performed on four replicates of each condition. 1ug of RNA was used for cDNA library preparation. Samples were sequenced using the Illumina HiSeqX platform to achieve 2 x 150 bp reads to generate 60 million paired-end reads per sample. FastqQC was performed as described in ([FastQC](https://www.bioinformatics.babraham.ac.uk/)) and read with Phred scores higher than 30 were aligned using HISAT2 (*62*). Feature counts function of the R Subread package was used to quantify the number of reads per transcript. Differential expression analysis was performed using DESeq2 (*63*). A log2FoldChange cutoff of 0.58 and P-value < 0.05 was used to identify DEGs. GO analysis was performed using ShinyGO online software (*64*).

### *In vitro* primary neuronal cultures

Neuronal cultures were performed from E15.5 embryos of *Lhx2^lox/lox^* and *NexCre/+; Lhx2^lox/lox^* mice. Embryos were dissected in ice-cold L-15 medium (Thermo Fischer Scientific SKU #41300039). pCAGGS-IRES-Cre-TdTomato + CAAX-EGFP (Progenitor mutant) and pCAGGS-IRES-TdTomato + CAAX-EGFP (Control) were electroporated in the dissected brains of *Lhx2^lox/lox^* mice. For the postmitotic loss condition, CAAX-EGFP was electroporated in *NexCre/+; Lhx2^lox/lox^* (postmitotic mutant) and littermate *Lhx2^lox/lox^* controls. The electroporated cortical explants were maintained on a Millicell Insert (Millipore, catalogue no. 217 PICM03050) with Neurobasal medium (ThermoFisher Scientific SKU #21103049) containing B-27 supplement (ThermoFisher Scientific SKU #17504044) and 1% GlutaMAX (ThermoFisher Scientific SKU #35050061) for 4 hours at 37°C with 5% CO_2_. The explants were dissociated into a single-cell suspension using 0.25%Trypsin (ThermoFisher Scientific SKU #15400054) and then dissociated by trituration after washes. The dissociated cells were plated on Poly-D-Lysine (Sigma catalogue no. P7280) coated coverslips in Neurobasal medium containing B-27 supplement, Penicillin/Streptomycin and 1% GlutaMAX for 15 days at a 5% CO_2_ atmosphere.

Dendritic morphology was analysed using the NeuronJ plugin in ImageJ. Apical and basal dendrites were identified based on their pyramidal morphology. Only neurons with just one apical dendrite were considered for analysis.

### *In vitro* patch-clamp electrophysiology

#### Slice preparation for in vitro patch-clamp recording

Mice were anaesthetised by intraperitoneal injection of a ketamine-xylazine mixture. After the onset of deep anaesthesia, assessed by cessation of the toe-pinch reflex, transcardial perfusion of the ice-cold cutting solution was performed. The cutting solution contained (in mM) 2.5 KCl, 1.25 NaH_2_PO_4_, 25 NaHCO_3_, 0.5 CaCl_2_, 7 MgCl_2_, 7 dextrose, 3 sodium pyruvate, and 200 sucrose (pH 7.3, ∼300 mOsm) saturated with carbogen (95% O_2_, 5% CO_2_). Thereafter, the brain was removed quickly, and 350-µm thick near-horizontal slices were prepared with a vibrating blade microtome (Leica Vibratome) while submerged in an ice-cold cutting solution saturated with carbogen. The slices were then incubated for 10–15 min at 34° C in a holding chamber containing a holding solution (pH 7.3, ∼300 mOsm) with the composition of (in mM) 125 NaCl, 2.5 KCl, 1.25 NaH_2_PO_4_, 25 NaHCO_3_, 2 CaCl_2_, 2 MgCl_2_, 10 dextrose, and 3 sodium pyruvate saturated with carbogen. Thereafter, the slices were kept in the holding chamber at room temperature for at least 30 minutes before recordings started.

#### Whole-cell current-clamp recordings

For electrophysiological recordings, slices were transferred to the recording chamber and were continuously perfused with carbogenated artificial cerebrospinal fluid (ACSF-extracellular recording solution) at a flow rate of 2–3 mL/min. All neuronal recordings were performed under current-clamp configuration at physiological temperatures (32–35° C) achieved through an inline heater that was part of a closed-loop temperature control system (Harvard Apparatus). The ACSF contained (in mM) 125 NaCl, 3 KCl, 1.25 NaH_2_PO_4_, 25 NaHCO_3_, 2 CaCl_2_, 1 MgCl_2_, and 10 dextrose (pH 7.3; ∼300 mOsm). Slices were first visualised under a 10× objective lens to visually locate layer II/III of the somatosensory cortex (S1). A 63× water-immersion objective lens was employed to perform visually guided patch-clamp recordings from mCherry-labelled pyramidal neurons (Pm control and Pm *Lhx2* neurons; unlabelled for Pg control and Pg *Lhx2* neurons) in superficial layers of S1 through a Dodt contrast microscope (Carl Zeiss Axioexaminer). Whole-cell current-clamp recordings were performed from layer II/III S1 pyramidal neurons with a Dagan BVC-700A amplifier.

Borosilicate glass electrodes with electrode tip resistance between 3 and 7 MΩ were pulled (P-97 Flaming/Brown micropipette puller; Sutter) from thick glass capillaries (1.5-mm outer diameter and 0.86-mm inner diameter; Sutter) and used for patch-clamp recordings. The pipette solution contained (in mM) 120 K-gluconate, 20 KCl, 10 HEPES, 4 NaCl, 4 Mg-ATP, 0.3 Na-GTP, and 7 K_2_-phosphocreatine (pH 7.3 adjusted with KOH; osmolarity ∼300 mOsm). Series resistance was monitored and compensated online with the bridge-balance circuit of the amplifier. Experiments were discarded only if the initial resting membrane potential was more depolarised than –60 mV if series resistance rose above 50 MΩ, or if there were fluctuations in temperature and ACSF flow rate during the experiment. Voltages have not been corrected for the liquid junction potential, which was experimentally measured to be ∼8 mV. Neuronal response to a 50-pA hyperpolarising current pulse was continuously monitored to observe and correct series resistance changes using the bridge balance circuit throughout the course of the experiment.

#### Pharmacological blockers

All recordings were performed in the presence of synaptic receptor blockers in the ACSF. Drugs and their concentrations used in the experiments were 10 µM 6-cyano-7-nitroquinoxaline-2,3-dione (CNQX), an AMPA receptor blocker; 10 µM (+) bicuculline and 10 µM picrotoxin, both GABA_A_ receptor blockers, and 2 µM CGP55845, a GABA_B_ receptor blocker (all synaptic blockers from Abcam) in the ACSF.

#### Subthreshold measurements

We characterised S1 pyramidal neurons with several electrophysiological measurements using standard protocols(*58–60*, *65*, *66*), detailed below. Resting membrane potential (*V*_*RMP*_) was measured as the voltage at which the cell rested when no current was injected (Supplementary Fig. S3–S4). Input resistance (*R*_*in*_) was measured as the slope of a linear fit to the steady-state voltage-current (V-I) plot obtained by injecting subthreshold current pulses of amplitudes spanning –50 to 0 pA in steps of 10 pA (Figure 2A–D). The sag ratio was measured from the voltage response of the cell to a hyperpolarising current pulse (Supplementary Fig. S3–S4). Sag ratio was defined as (*V*_*initial*_/*V*_*SS*_), where *V*_*SS*_ and *V*_*initial*_ depict the steady-state and peak (during the initial 50-ms period after current injection) voltage deflections (from *V*_*RMP*_), respectively. To assess temporal summation, five alpha excitatory postsynaptic potentials (*α*-EPSPs) with 50-ms intervals were injected as currents of the form *I*_*α*_= *I*_*max*_*t exp*(−*αt*), with *α* = 0.1 ms^−1^. The temporal summation ratio (*S*_*α*_) in this train of five EPSPs (Supplementary Fig. S3–S4) was computed as *E*_*last*_/*E*_*first*_, where *E*_*last*_ and *E*_*first*_ were the amplitudes of the last and first EPSPs in the train, respectively.

The chirp stimulus, a sinusoidal current with its frequency linearly spanning 0–15 Hz in 15 s and of constant amplitude adjusted to be below the firing threshold, was used to characterise the impedance profiles (Supplementary Fig. S3–S4). The voltage response of the neuron was recorded for chirp current stimulus injection at *V*_*RMP*_. The ratio of the Fourier transform of the voltage response to the Fourier transform of the chirp stimulus formed the impedance profile. The frequency at which the impedance amplitude reached its maximum was the resonance frequency (*f*_*R*_). Resonance strength (*Q*) was measured as the ratio of the maximum impedance amplitude to the impedance amplitude at 0.5 Hz. The total inductive phase (*Ф*_*L*_) was defined as the area under the inductive part of the impedance phase profile as a function of frequency (Supplementary Fig. S3–S4).

#### Suprathreshold measurements

Action potential (AP) firing frequency was computed by extrapolating the number of spikes obtained during a 700-ms current injection to 1s. The amplitude of these pulse current injections was varied from 0 pA to 250 pA in steps of 50 pA to construct the firing frequency *vs* injected current (*f–I*) plot (Figure 2E–H). In addition, to assess firing at higher current injections, we constructed *f–I* plots with high current injections ranging from 250–1250 pA in steps of 250 pA (Supplementary Fig. S5C–D). Various AP-related measurements(*58–60*, *65*, *66*) were derived from the first action potential in the voltage response of the cell, resting at *V*_*RMP*_, to a 500-pA pulse current injection (Supplementary Figures S6–S7). AP amplitude (*V*_*AP*_) was computed as the difference between the peak voltage of the spike (*V*_*peak*_) and *V*_*RMP*_(Supplementary Fig. S6–S7). The temporal distance between the timing of the first spike and the time of the current injection was defined as latency to the first spike *T*_1*AP*_ (Supplementary Fig. S6–S7). The duration between the first and the second spikes was defined as the first inter-spike interval (Supplementary Fig. S6–S7). AP half-width (*T*_*APHW*_) was the temporal width measured at the half-maximal points of the AP peak concerning *V*_*RMP*_. The maximum and minimum rate of change of voltage (*dV*/*dt*) of the AP temporal derivatives was calculated from the temporal first derivative of the voltage trace (Supplementary Fig. S6– S7). The voltage in the AP trace corresponds to the timepoint at which the *dV*/*dt* crossed 20 V/s defined as the AP threshold *V*_*t*ℎ_ (Supplementary Fig. S6–S7).

#### Analyses of electrophysiological data and statistics

All data acquisition and analyses were performed with custom-written software in IGOR Pro (WaveMetrics), and statistical analyses were performed using the Wilcoxon rank sum test in the R computing package (http://www.r-project.org/).

