## Supplementary Figures for "LHX2 regulates dendritic morphogenesis in layer II/III of the neocortex via distinct pathways in progenitors and postmitotic neurons"

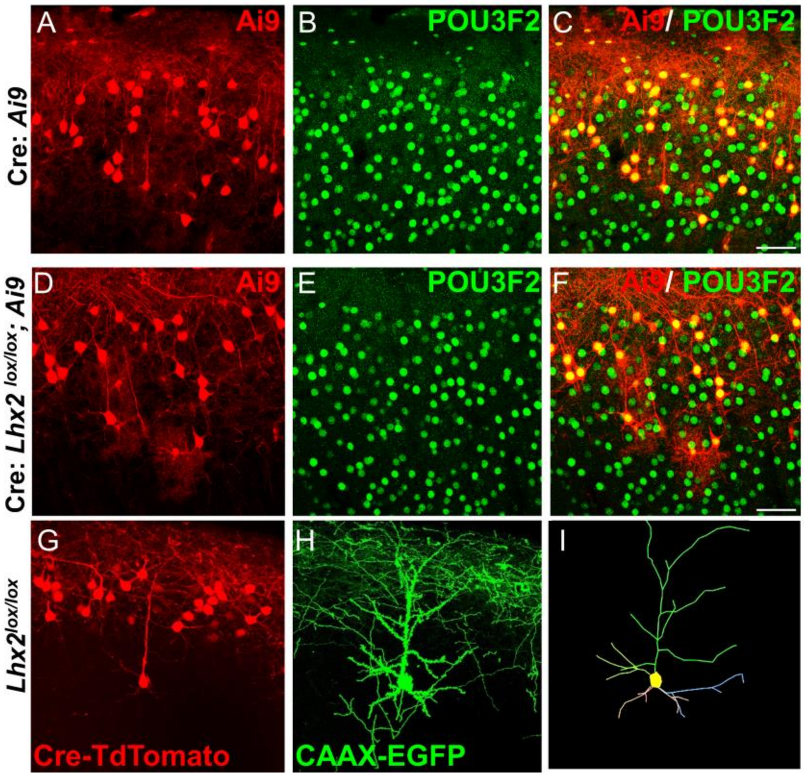


**Supplementary Figure S1: Loss of *Lhx2* from E15.5 progenitors affects dendritic arborisation of layer II/III neurons without affecting subtype identity.**

(A–F) Expression of upper-layer specific marker POU3F2 is not affected in *Lhx2* mutant neurons. Electroporation of *Cre* in *Ai9/+* at E15.5 (A) labels POU3F2-positive upper layer neurons (B, C). Loss of *Lhx2* by electroporation of *Cre* in *Lhx2*$^{lox/lox}$*; Ai9/+* progenitors at E15.5 (D) does not affect the expression of the upper-layer identity marker POU3F2 (E, F). (A–C) Example electroporated-neuron reconstruction at P30. Electroporation of Cre-TdTomato (A) creates cell-autonomous deletion in the progenitors of *Lhx2* condition knockout mice (*Lhx2*$^{lox/lox}$). Simultaneous membrane-bound CAAX-EGFP (B) electroporation leads to proper dendritic arbour visualisation. These arbours are 3D reconstructed using the Neurolucida software (C). All analyses were performed at P30. *Scale bars are 50 μm.*

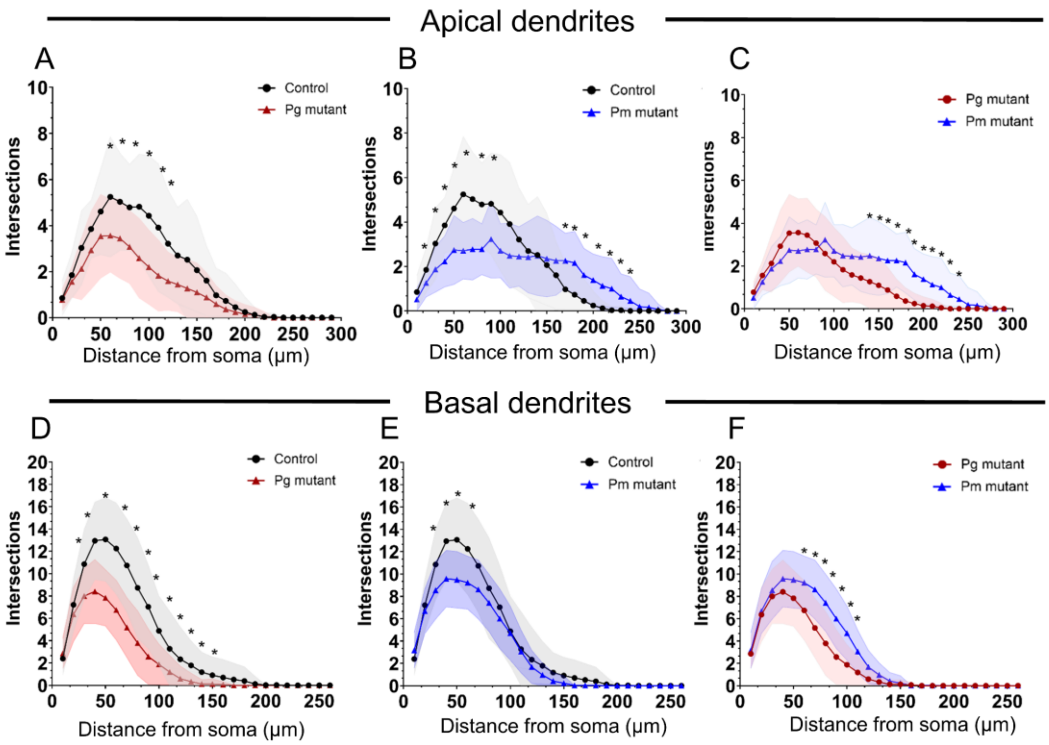


**Supplementary Figure S2: Progenitor and postmitotic loss of *Lhx2* lead to distinct levels of dendritic complexity defects.**

Pairwise comparisons between Control (black line), Pg mutant (red line), and Pm mutant (blue line) apical dendrite (A–C) and basal dendrite (D–F) complexities with asterisks marking significant differences as shown in Figure 1J, K. 35 to 40 neurons were scored for each condition obtained from 4 biologically independent replicates; *Statistical test: Multiple T Tests (two-tailed). *(p<0.05).*

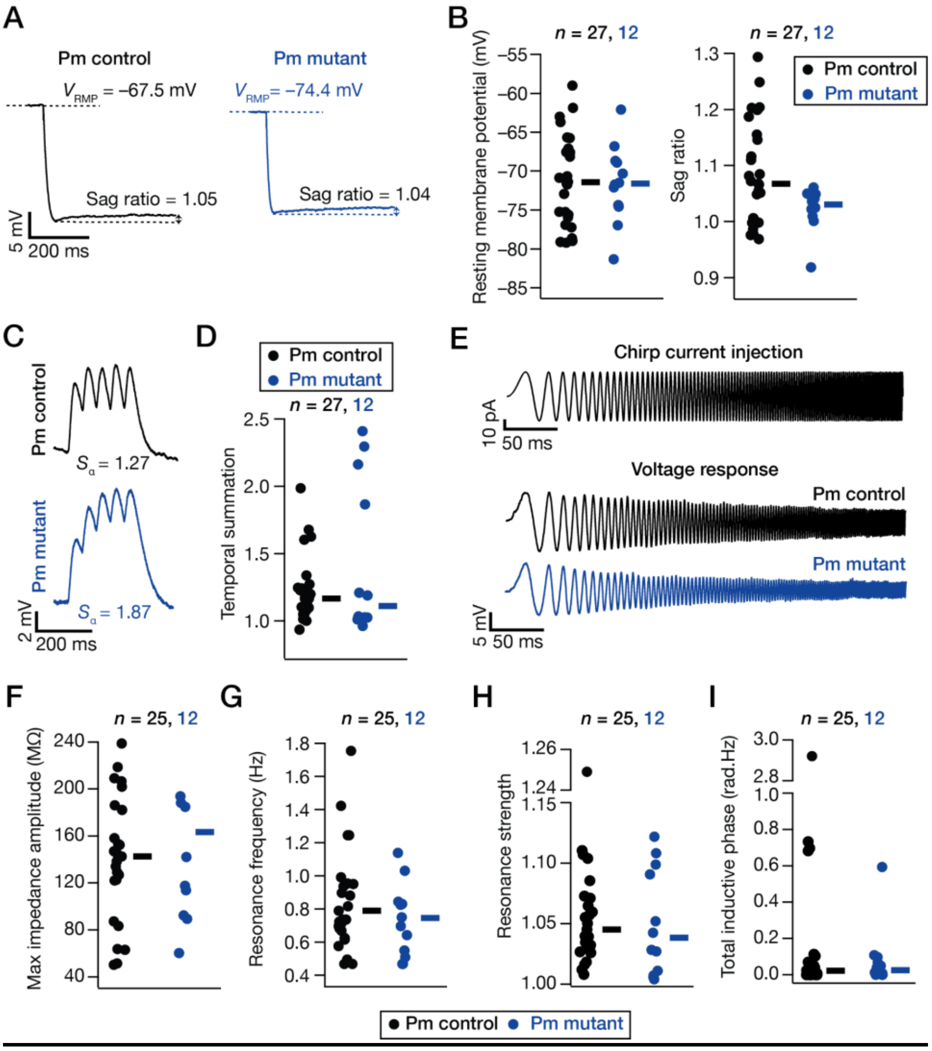


**Supplementary Figure S3: Sub-threshold measurements for neocortical layer II/III pyramidal neurons from control and postmitotic (Pm) *Lhx2* mutant groups.**

(A) Traces showing voltage response to a 250-pA hyperpolarising pulse current for the example control (left) and Pm mutant (right) neurons. The resting membrane potential ($V_{RMP}$) is shown with a dotted line on top of the trace just before current injection. Sag is represented as the difference between the initial peak and the steady-state voltage deflection. (B) Beeswarm plots of resting membrane potential ($V_{RMP}$) (left) and Sag ratio (right) recorded from all control and Pm mutant neurons. (C) Train of five excitatory postsynaptic potentials ($\alpha$-EPSPs) recorded from example control (top) and Pm mutant (bottom) neurons. Temporal summation ($S_\alpha$) was calculated as a ratio between the last and the first EPSP amplitudes. (D) Beeswarm of $S_\alpha$ recorded from all control and Pm mutant neurons. (E) *Top*, Chirp current stimulus. *Bottom*, Voltage responses of the example control and mutant neurons to the chirp current stimulus. (F–I) Beeswarm plots of maximum impedance amplitude (F), resonance frequency, $f_R$ (G), resonance strength, $Q$ (H), and total inductive phase, $\phi_L$ (I) calculated for all control and Pm mutant neurons. *Statistical test: Wilcoxon rank sum test. n represents the number of total neurons recorded in each group. The thick line on the right of each beeswarm plot depicts respective median values.*

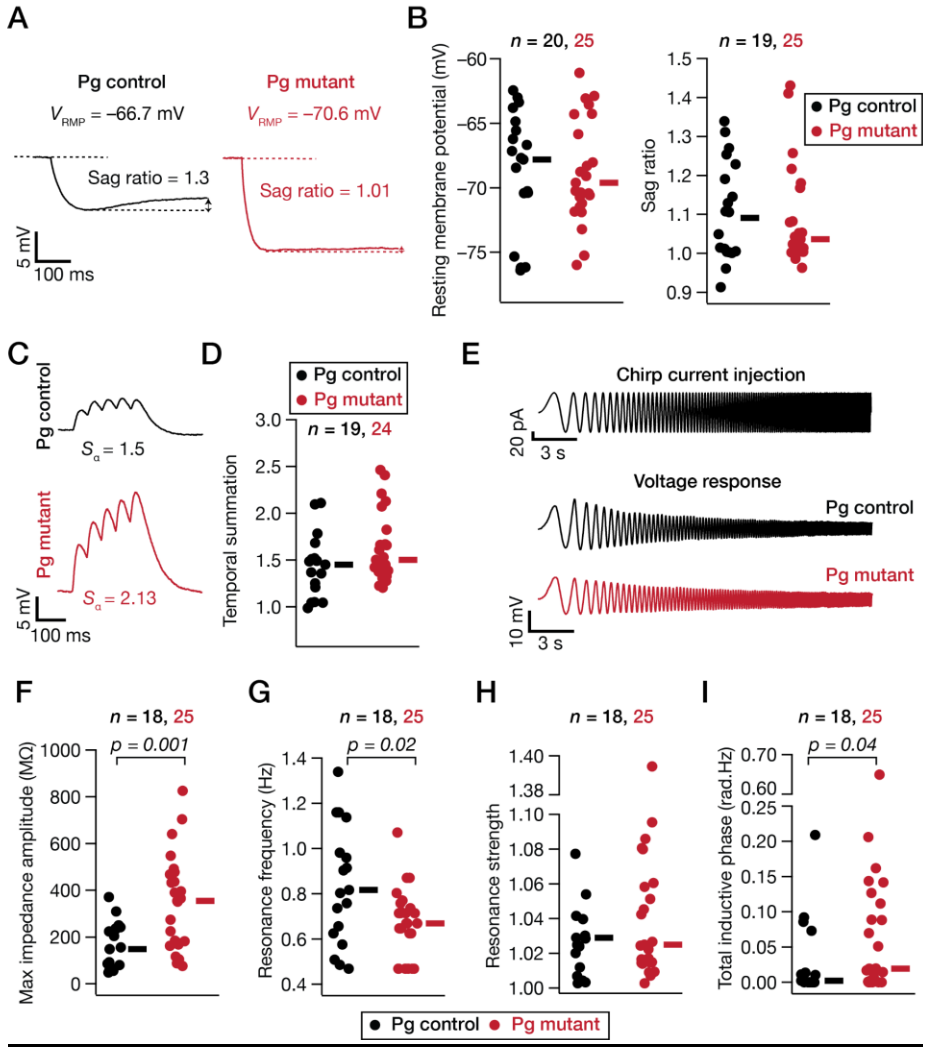


**Supplementary Figure S4: Sub-threshold measurements for neocortical layer II/III pyramidal neurons from control and progenitor (Pg) *Lhx2* mutant groups.**

(A) Traces showing voltage response to a 50-pA hyperpolarising pulse current for the example control (left) and Pg mutant (right) neurons. The resting membrane potential ($V_{RMP}$) is shown with dotted line on top of the trace just before current injection. Sag is represented as the difference between the initial peak and the steady-state voltage deflection. (B) Beeswarm plots of resting membrane potential ($V_{RMP}$) (left) and Sag ratio (right) recorded from all control and Pg mutant neurons. (C) Train of five excitatory postsynaptic potentials ($\alpha$-EPSPs) recorded from example control (top) and Pg mutant (bottom) neurons. Temporal summation ($S_\alpha$) was calculated as ratio between the last and the first EPSP amplitudes. (D) Beeswarm of $S_\alpha$ recorded from all control and Pg mutant neurons. (E) *Top*, Chirp current stimulus. *Bottom*, Voltage responses of the example control and mutant neurons to the chirp current stimulus. (F–I) Beeswarm plots of maximum impedance amplitude (F), resonance frequency, $f_R$ (G), resonance strength, $Q$ (H), and total inductive phase, $\phi_L$ (I) calculated for all control and Pg mutant neurons. *Statistical test: Wilcoxon rank sum test. n represents the number of total neurons recorded from in each group. The thick line on the right of each beeswarm plot depicts respective median values.*

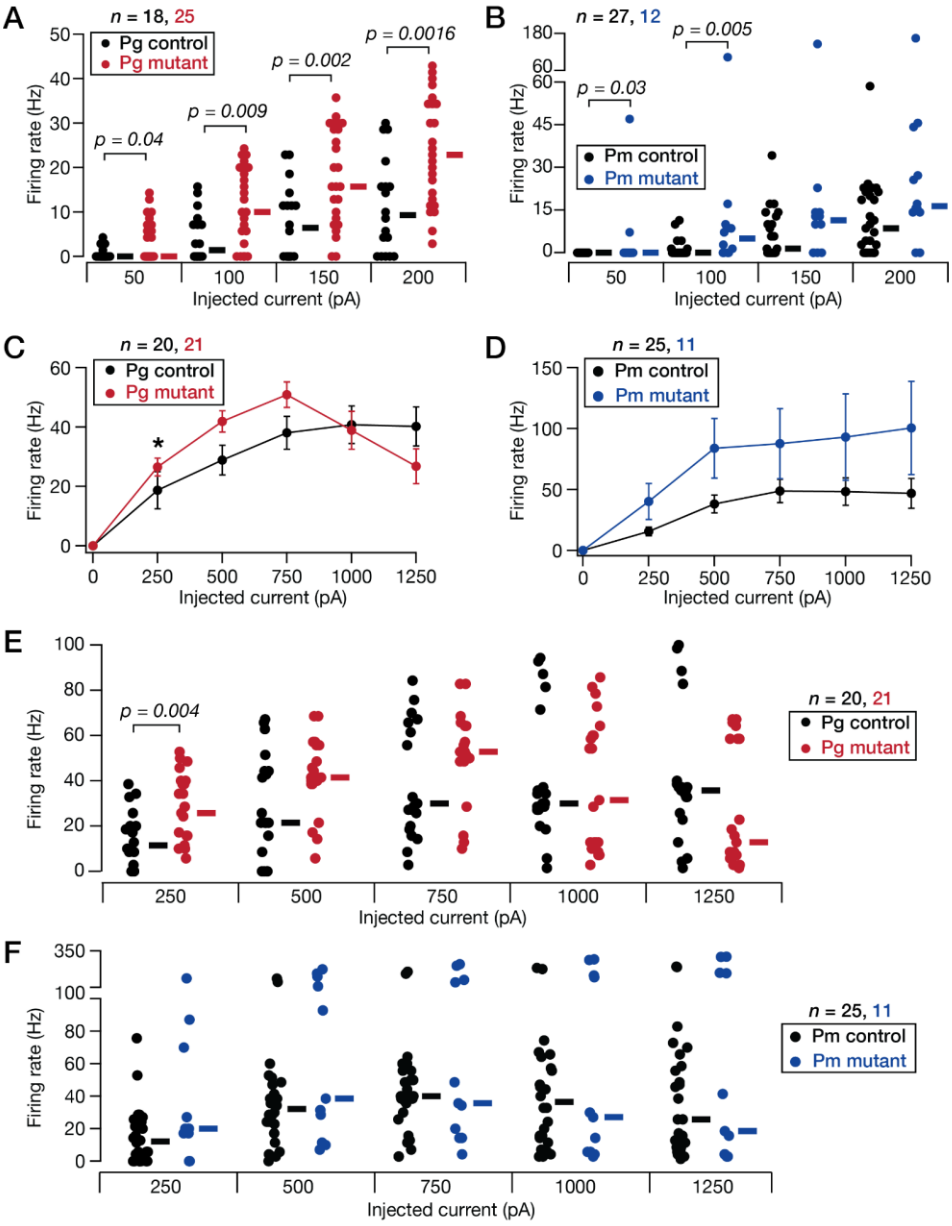


**Supplementary Figure S5: Loss of *Lhx2* in progenitors (Pg) or postmitotic (Pm) neurons enhanced intrinsic excitability of neocortical layer II/III pyramidal neurons.**

(A) Beeswarm plots of action potential firing frequencies for current injection amplitudes from 50–200 pA for control and Pg mutant neurons. (B) Same as panel A, for Pm mutant neurons and associated controls. (C) Mean-SEM plot showing firing rates of neurons from control and Pg mutant mice, plotted against current injection amplitudes from 250–1250 pA. (D) Same as panel C, for Pm mutant neurons and associated controls. (E) Beeswarm plots of action potential firing frequencies for current injection amplitudes from 250–1250 pA for control and Pg mutant neurons. (F) Same as panel E, for Pm mutant neurons and associated controls. *Statistical test: Wilcoxon rank sum test. n represents the number of total neurons recorded in each group. The thick line on the right of each beeswarm plot depicts respective median values.*

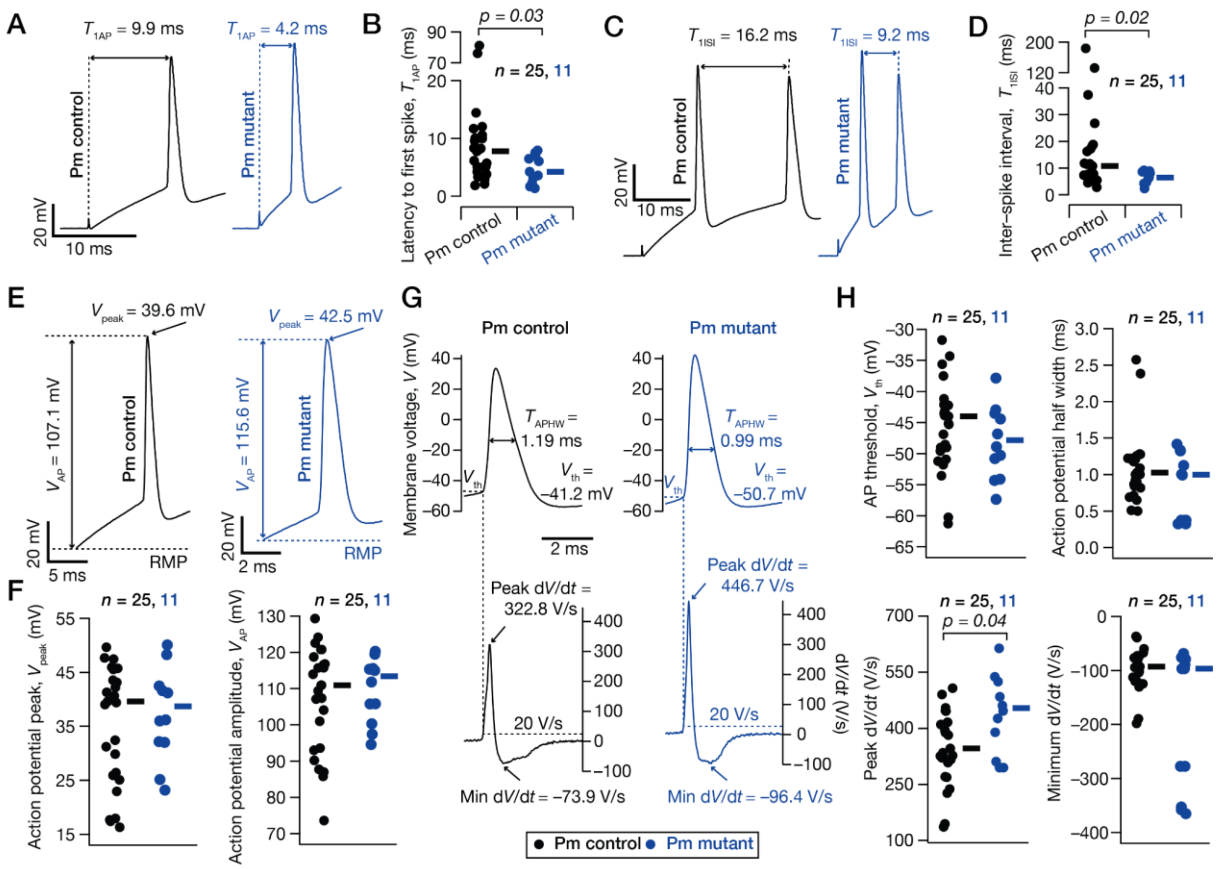

**Supplementary Figure S6: Supra-threshold measurements for neocortical layer II/III pyramidal neurons from control and postmitotic (Pm) *Lhx2* mutant groups.**

(A) The first action potential in a spike train, recorded in response to a 500-pA current injection into example neurons from control (left) and Pm mutant (right) groups. Arrows mark the latency to first spike ($T_{1AP}$), measured from the start time of current injection. (B) Beeswarm plot of $T_{1AP}$ recorded from all control and Pm mutant neurons. (C) The first two action potentials in a spike train, recorded in response to a 500-pA current injection into example neurons from control (left) and Pm mutant (right) groups. Arrows mark the first inter-spike interval ($T_{ISI}$). (D) Beeswarm plot of $T_{ISI}$ recorded from all control and Pm mutant neurons. (E) The first action potential in a spike train, recorded in response to a 500-pA current injection into example neurons from control (left) and Pm mutant (right) groups. Top arrow marks the peak voltage deflection $V_{peak}$ . $V_{AP}$ marks the action potential amplitude, measured from resting potential. (F) Beeswarm plots of $V_{peak}$ and $V_{AP}$ for control and Pm mutant groups. (G) The first action potential (top) in a spike train, recorded in response to a 500-pA current injection into example neurons from control (left) and Pm mutant (right) groups. The respective temporal derivatives ($dV/dt$) are shown below. Arrows mark the peak and the minimum $dV/dt$ associated with each action potential. (H) Beeswarm plots of action potential threshold ($V_{th}$), action potential half-width ($T_{APHW}$), peak $dV/dt$, and minimum $dV/dt$ from control and Pm mutant neurons. *Statistical test: Wilcoxon rank sum test. n represents the number of total neurons recorded in each group. The thick line on the right of each beeswarm plot depicts respective median values.*

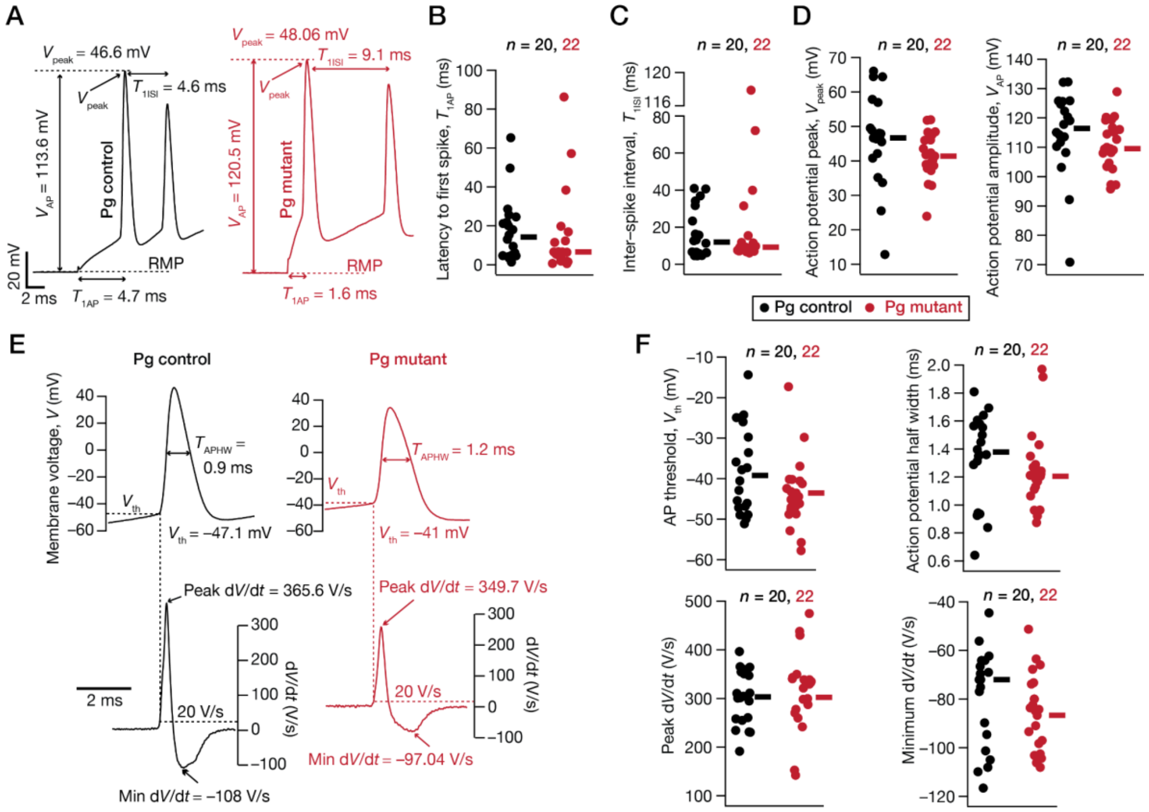


**Supplementary Figure S7: Supra-threshold measurements for neocortical layer II/III pyramidal neurons from control and progenitor (Pg) *Lhx2* mutant groups.**

(A) The first two action potentials in a spike train, recorded in response to a 500-pA current injection into example neurons from control (left) and Pg mutant (right) groups. Arrows mark the latency to first spike ($T_{1AP}$), measured from the start time of current injection, the first inter-spike interval ($T_{ISI}$). Top arrows mark the peak voltage deflection $V_{peak}$. $V_{AP}$ marks the action potential amplitude, measured from resting membrane potential (RMP). (B) Beeswarm plot of recorded from all control and Pg mutant neurons. (C) Beeswarm plot of $T_{ISI}$ recorded from all control and Pg mutant neurons. (D) Beeswarm plots of $V_{peak}$ and $V_{AP}$ for control and Pg mutant groups. (E) The first action potential (top) in a spike train, recorded in response to a 500-pA current injection into example neurons from control (left) and Pg mutant (right) groups. The respective temporal derivatives ($dV/dt$) are shown below. Arrows mark the peak and the minimum $dV/dt$ associated with each action potential. (F) Beeswarm plots of action potential threshold ($V_{th}$), action potential half-width ($T_{APHW}$), peak $dV/dt$, and minimum $dV/dt$ from control and Pg mutant neurons. *Statistical test: Wilcoxon rank sum test. n represents the number of total neurons recorded in each group. The thick line on the right of each beeswarm plot depicts respective median values.*

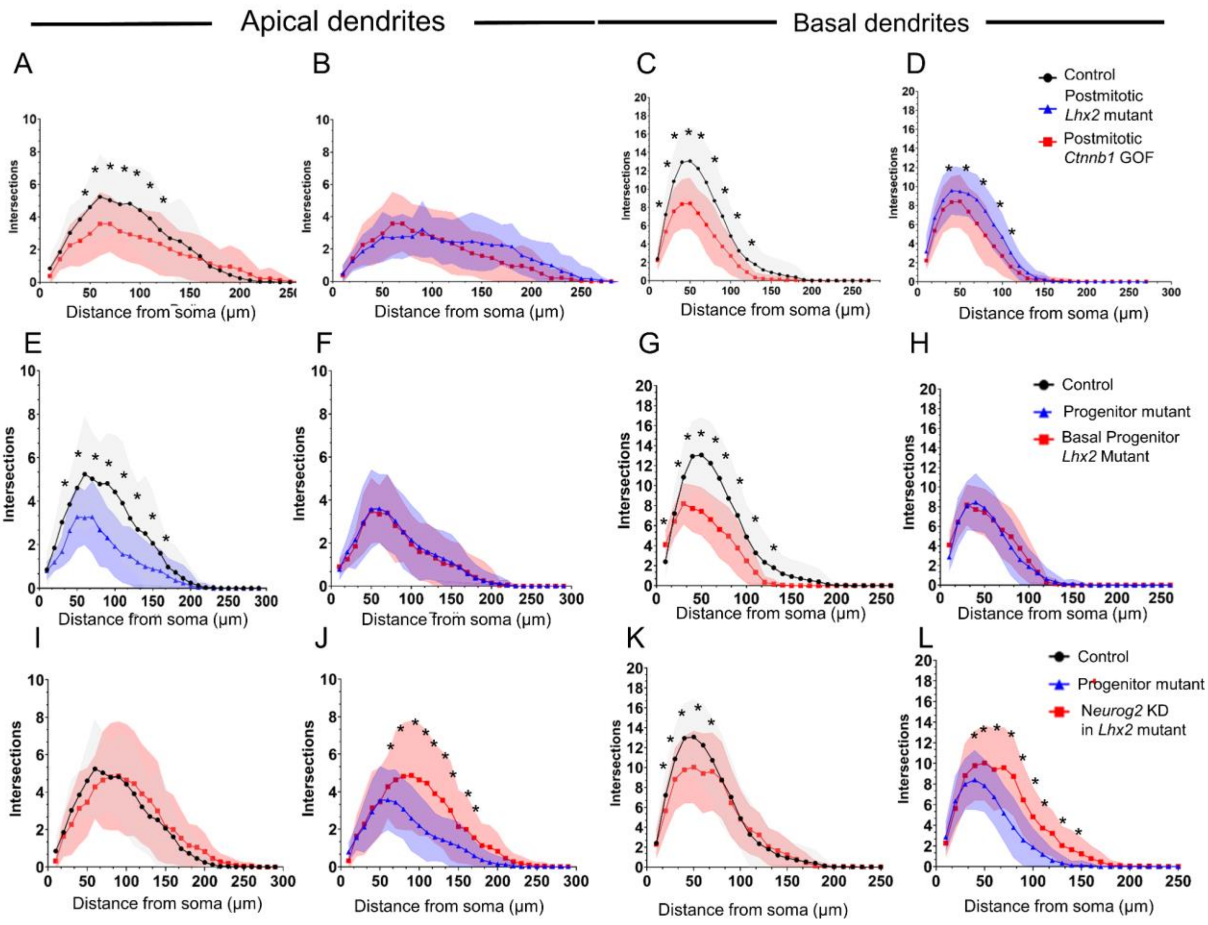


**Supplementary Figure S8: Individual comparison of Sholl analysis from different conditions.**

(A–D) Pairwise comparisons of apical (A, B) and basal (C, D) from Control, Pm mutant and *Ctnnb1* GOF conditions as represented in Figure 3J, K. (E–H) Pairwise comparisons of apical (E, F) and basal (G, H) from Control, Pg mutant and *Eomes*-Cre driven basal progenitor mutant conditions as represented in Figure 4K, L. (E–H) Pairwise comparisons of apical (I, J) and basal (K, L) from Control, Pg mutant and *Neurog2* knockdown in *Lhx2* Pg mutant conditions as represented in Figure 4N, O. (30–40 neurons were analysed from 3 biologically independent experiments. *Statistical test: Multiple T Tests (two-tailed). *(p<0.05).*

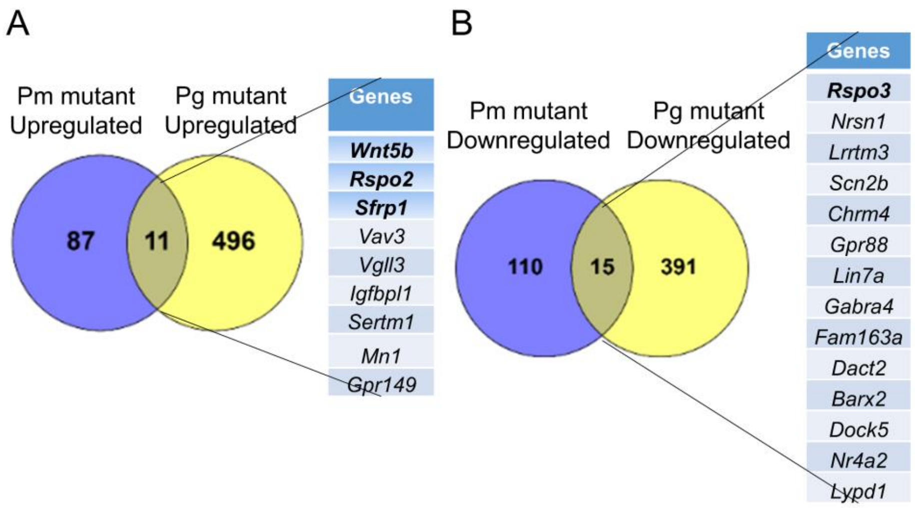


**Supplementary Figure S9: Comparison of transcriptomic changes in Pg versus Pm loss of *Lhx2*.**

List of common genes upregulated (A) or downregulated (B) upon progenitor (Pg) and postmitotic (Pm) deletion of *Lhx2* in layer II/III cortical neurons analyzed at P5. 583 genes are upregulated and 501 genes are downregulated genes that do not overlap in Pg and Pm mutants, whereas 11 upregulated and 15 downregulated genes overlap Pm mutant RNA Seq does not reveal an upregulation of *Neurog2.* Common genes between Pg and Pm RNA Seq include certain Wnt signalling genes like *Wnt5b, Rspo2, Rspo3 and Sfrp1,* among others.

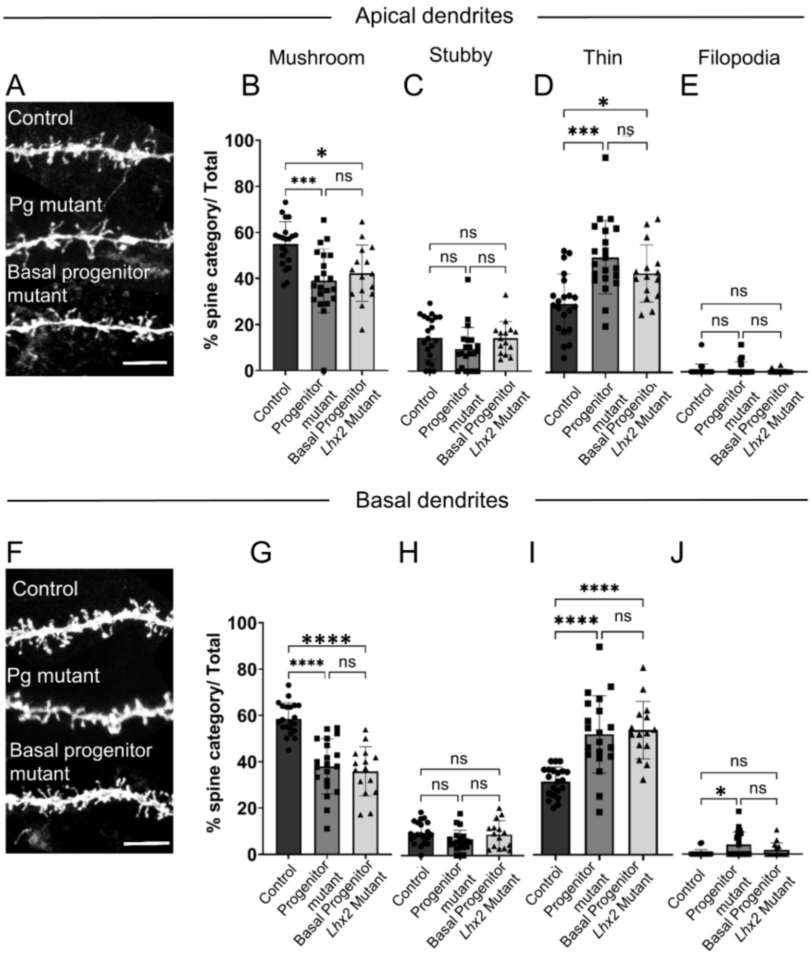


**Supplementary Figure S10: Basal progenitor specific *Lhx2* deletion recapitulates pan-progenitor specific spine defects.**

(A–J) Dendritic spine morphology is affected in *Eomes*-Cre driven basal progenitor *Lhx2* mutant neurons. In apical and basal dendrites, the percentage of spines with mushroom morphology was significantly reduced (B, G) with a concomitant increase in thin morphology (D, I), similar to progenitor (Pg) mutants. The percentage of stubby spines was, however, not affected (C, H). The percentage of filopodial spines increased in Pg basal dendrites. 20 dendrites with 25 spines each were analysed from each category. *Statistical Test: Kruskal-Wallis Test*. *Scale bar: 5µm.*

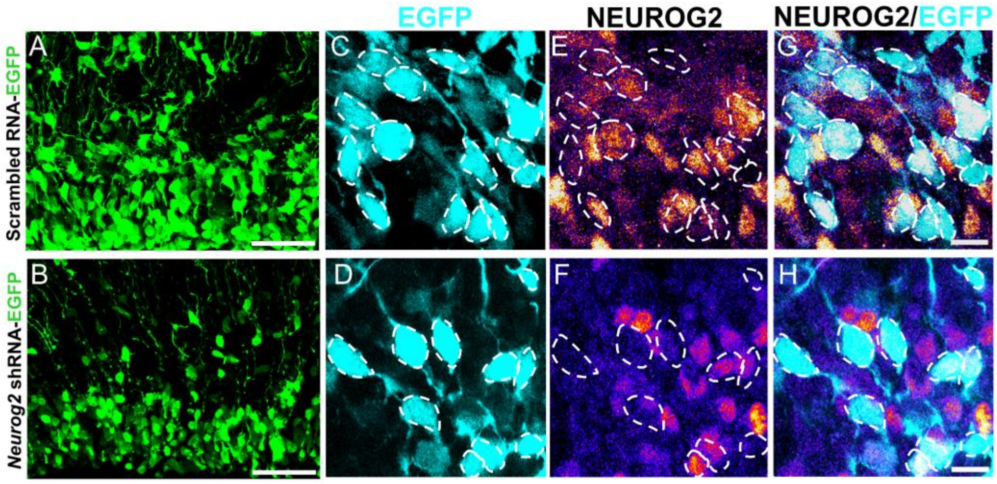


**Supplementary Figure S11: *Neurog2* shRNA plasmid electroporation downregulates NEUROG2 expression.**

(A, B) Electroporation of scrambled pSilcaggs-scrambled RNA-EGFP (A) and pSilcaggs-*Neurog2* shRNA-EGFP (B) at E15.5 and analysed at E16.5. Downregulation of *Neurog2* leads to delayed migration of cells (B), as previously described in (*44*). (C, D) *Neurog2* shRNA-EGFP (D) reduces NEUROG2 levels in electroporated cells compared to scrambled RNA (C). *N=2. Scale bars in (A, B) are 50μm, and (G, H) are 10μm.*

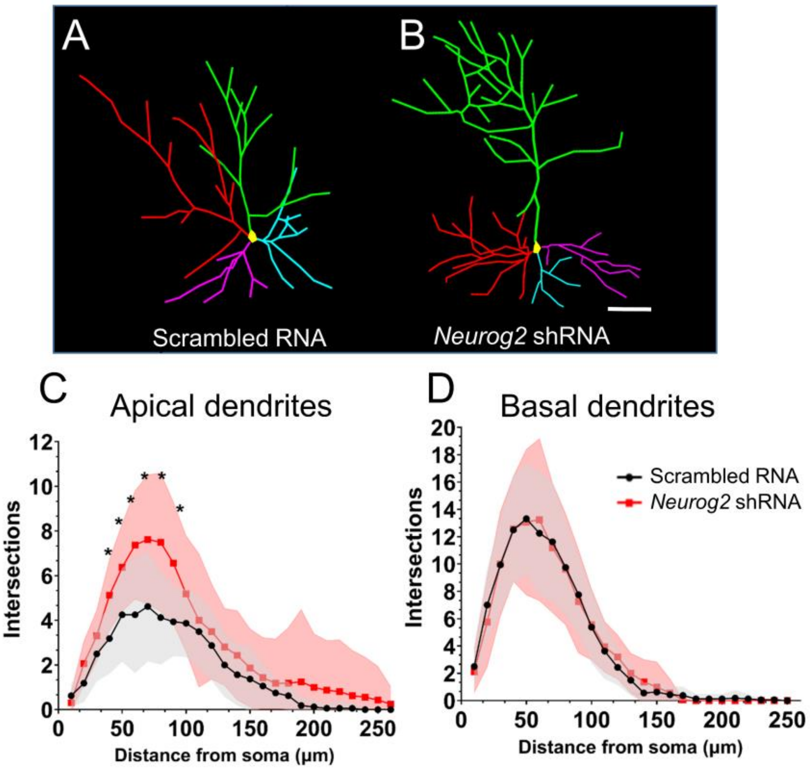


**Supplementary Figure S12: Downregulation of *Neurog2* in E15.5 progenitors results in increased dendritic complexity of the apical dendrites in Layer II/III neurons.**

(A, B) Neurolucida tracings of neurons labelled at E15.5 via *in utero* electroporation of CAAX-mCherry and analyzed at P25. The morphology of neurons co-electroporated with scrambled RNA (black line) compared with those co-electroporatedwith *Neurog2* shRNA (red line) displays a significant increase in the complexity of apical dendrites (C) upon *Neurog2* downregulation, whereas no significant changes are observed in the basal dendrites (D). 16 neurons were scored for each condition obtained from 2 biologically independent replicates. *Statistical test: Multiple t test., *p<0.05. Scale bar: 50 µm.*

### Supplementary Tables:

**Supplementary Data S1**: Statistical correlation of Sholl analysis for different datasets.

**Supplementary Data S2**: P5 Control versus Progenitor *Lhx2* mutant neuron RNA Seq differentially expressed gene list.

**Supplementary Data S3**: P5 Control versus Postmitotic *Lhx2* mutant neuron RNA Seq differentially expressed gene list.
